# Neuroinflammation and EIF2 signaling persist in an HiPSC tri-culture model of HIV infection despite antiretroviral treatment

**DOI:** 10.1101/779025

**Authors:** Sean K. Ryan, Michael V. Gonzalez, James P. Garifallou, Frederick C. Bennett, Kimberly S. Williams, Hakon Hakonarson, Stewart A. Anderson, Kelly L. Jordan-Sciutto

## Abstract

HIV-Associated Neurocognitive Disorders (HAND) affect over half of HIV-infected individuals worldwide, despite antiretroviral therapy (ART). Therapeutically targetable mechanisms underlying HAND remain elusive. We developed a human-induced pluripotent stem cell (HiPSC) based model; whereby, we independently differentiate HiPSCs into neurons, astrocytes, and microglia and systematically combine to generate a tri-culture with or without HIV-infection and ART. scRNAseq analysis on tri-cultures including HIV-infected microglia revealed inflammatory signatures in the microglia and EIF2 signaling in all three cell types. Remarkably, EFZ alone induced a similar response to infection. Treatment with the antiretroviral compound Efavirenz (EFZ) mostly resolved these signatures; However, EFZ increased RhoGDI and CD40 signaling in the HIV-infected microglia. This activation was associated with a persistent increase in TNFa expression. This work establishes a tri-culture that recapitulates key features of HIV infection in the CNS and provides a new model to examine the effects of HIV infection and its treatment with antiretrovirals.

## Introduction

HIV-Associated Neurocognitive Disorders (HAND) are a chronic, progressing spectrum of diseases that lead to a range of neurologic disorders, including HIV-associated dementia. The major pathologic manifestation that persists in HAND patients with antiretroviral treatment (ART) is synaptodendritic damage and the accumulation of microglia, the immune cell in the central nervous system (CNS); however, the mechanisms underlying synaptic damage remain elusive. Synapse loss is associated with infiltration of macrophages from outside of the CNS and activation of microglia. Both HIV-infected macrophage populations can release cytokines, viral proteins, and excitotoxins, which can lead to synaptic damage (Saylor et al., 2016). Activated microglia aberrantly phagocytosing synapses have been described in multiple neurological disorders including: Alzheimer’s, Frontotemporal Dementia, and Multiple Sclerosis (Keren-Shaul et al., 2017; Lui et al., 2016; Salter & Stevens, 2017; Zrzavy et al., 2017). While patients on ART experience milder forms of HAND, they still experience chronic inflammation (Kolson, 2017). Infected microglia and uninfected, but active microglia may be working in tandem to slowly release proinflammatory cytokines and reactive oxygen species that, over time, can lead to synaptodendritic damage.

Inflammation, via various pathways, plays a large role in the neurodegeneration seen in HAND. Our lab and others have implicated the unfolded protein response (UPR), a part of the integrated stress response (ISR) in neurodegeneration including in HAND (Walter & Ron, 2011). EIF2 activation, downstream of UPR activation, has been implicated in HAND (C. Akay et al., 2012; K. A. Lindl, Akay, Wang, White, & Jordan-Sciutto, 2007), progressive supranuclear palsy and Alzheimer’s disease (Stutzbach et al., 2013). CD40 has also been implicated in viral replication control and microglia activation during HIV infection (D’Aversa, Eugenin, & Berman, 2008; Kornbluth, 2000; Sui et al., 2007). CD40 signaling is implicated in multiple cellular processes including inflammatory response. Specifically, CD40 activation can lead to, CD40 activation can lead to activation of the NF-κB non canonical pathway and the production of IL-8 and TNFa (D’Aversa et al., 2008; Homig-Holzel et al., 2008). TNFa, in turn, can activate the NADPH oxidase complex (Y.-S. Kim, Morgan, Choksi, & Liu, 2007). Similarly, RhoGDI normally anchors Rac1, but during inflammation, Rac1 is released from RhoGDI and moves to the plasma membrane where it interacts with the NADPH oxidase complex which creates superoxide when activated (Wilkinson & Landreth, 2006) and regulates cytokine and neurotoxin release by microglia exposed to HIV-Tat (Turchan–Cholewo et al., 2009). Understanding how various endogenous pathways may influence HAND using a therapeutic relevant model is important.

While the major pathological manifestation of HAND is synaptodendritic damage, the response during initial exposure to HIV and ART is unknown partly because current models for HIV-mediated neuroinflammation are limited by species differences and human tropism of the virus. For instance, the HIV-1 Tat transgenic mouse model exhibits neuroinflammation and behavioral deficits, but only expresses a single viral protein from astrocytes (Carey, Sypek, Singh, Kaufman, & McLaughlin, 2012; B. O. Kim et al., 2003; Langford et al., 2018). Another HIV-1 transgenic mouse model has neuroinflammation, but is missing two of the genes, *gag* and *pol*, and is expressed in every cell in the body (Reid et al., 2001). Lastly, while isolated microglia from human patients can give great insight, it is only a snapshot of the end stage of the disease. Therefore, the development of a novel *in vitro* human system allowing the interaction of the main cell types involved in HAND is needed to further understand the neuropathogenesis and develop novel therapeutics.

We have developed a Human-induced pluripotent stem cell (HiPSC) based model whereby we separately differentiate HiPSCs into forebrain-like excitatory neurons, astrocytes, and microglia then combine them to create a tri-culture, with or without HIV-infection of the microglia, and with or without ART. Our protocol rapidly produces microglia-like cells (iMg) that express multiple classical markers in mono-culture, productively infect with HIV, and respond to ART. In addition, we have developed a differentiation protocol for astrocyte-like cells (iAst) that express hallmark proteins. Utilizing this system, we investigated the effects of Infection (Inf), Infection with the antiretroviral Efavirenz (EFZ) (Inf+EFZ), and EFZ treatment alone (Uninf+EFZ), compared to uninfected tri-cultures (Uninf) and to each other. We found that EFZ treatment in infected conditions did not simply attenuate the inflammatory response but created a distinct reaction. In addition, EFZ treatment alone invoked its own discrete inflammatory response. We revealed gene expression changes leading to activation of EIF2 signaling in iMg, iAst, and iNrn, and highly activated inflammatory cytokine signaling in iMg in the Inf condition. EIF2 signaling persisted exclusively in the iNrns in the Inf+EFZ condition. While infection lead to increased inflammation in the iMg, it also led to decreased synaptophagy. EFZ treatment in infected cultures did not reverse the phagocytosis impairment but did quell the inflammation except for distinct gene activation of CD40 and RhoGDI signaling in the iMg. Lastly EFZ treatment itself lead to a very similar response as Inf. Our system has reveals the complex roles of the individual cell types during infection ± ART and how ART alone can elicit an inflammatory response.

## Results

### iMicroglia exhibit similar gene and protein expression as other iPSC-derived microglia

We modified two previously published protocols to generate the iMg (Abud et al., 2017; Paluru et al., 2014) (Figure 1a). The first step is a nine-day differentiation of iPSCs into CD41+CD235+ common myeloid progenitors (CMPs) (Paluru et al., 2014). The second step is a modified 11-day protocol (Abud et al., 2017) that produces a highly pure population of iMg. We tested this protocol on four separate cell lines. These iMg show ramification (Figure 1-figure supplement 1a), express several classical markers, including: CX3CR1, IBA1, and P2RY12, at the protein and RNA level (Figures 1b and 1c) in mono-culture, and have a significantly different gene expression profile than monocyte-derived macrophages (MDMs) (Figure 1-figure supplement 1b). We validated the findings in BulkRNAseq with qRT-PCR. The iMg exhibited a 179-fold, 28-fold, and 11-fold increase in *CX3CR1*, *P2RY12*, and *TMEM119* respectively over MDMs (Figure 1d-1f). Importantly they also express CCR5 (Figure 1-figure supplement 1c), one of the co-receptors necessary for HIV infection (Deng et al., 1996). Utilizing PCA, we compared our iMg to human MDMs we generated and the primary human adult and fetal microglia, iPSCs, induced hematopoietic progenitor cells, and iMicroglia from (Abud et al., 2017). PCA revealed considerate clustering of our iMg (pink) to the primary human microglia (blue) and previously published iPSC-derived microglia (light orange) (Figure 1g).

**Figure 1:**
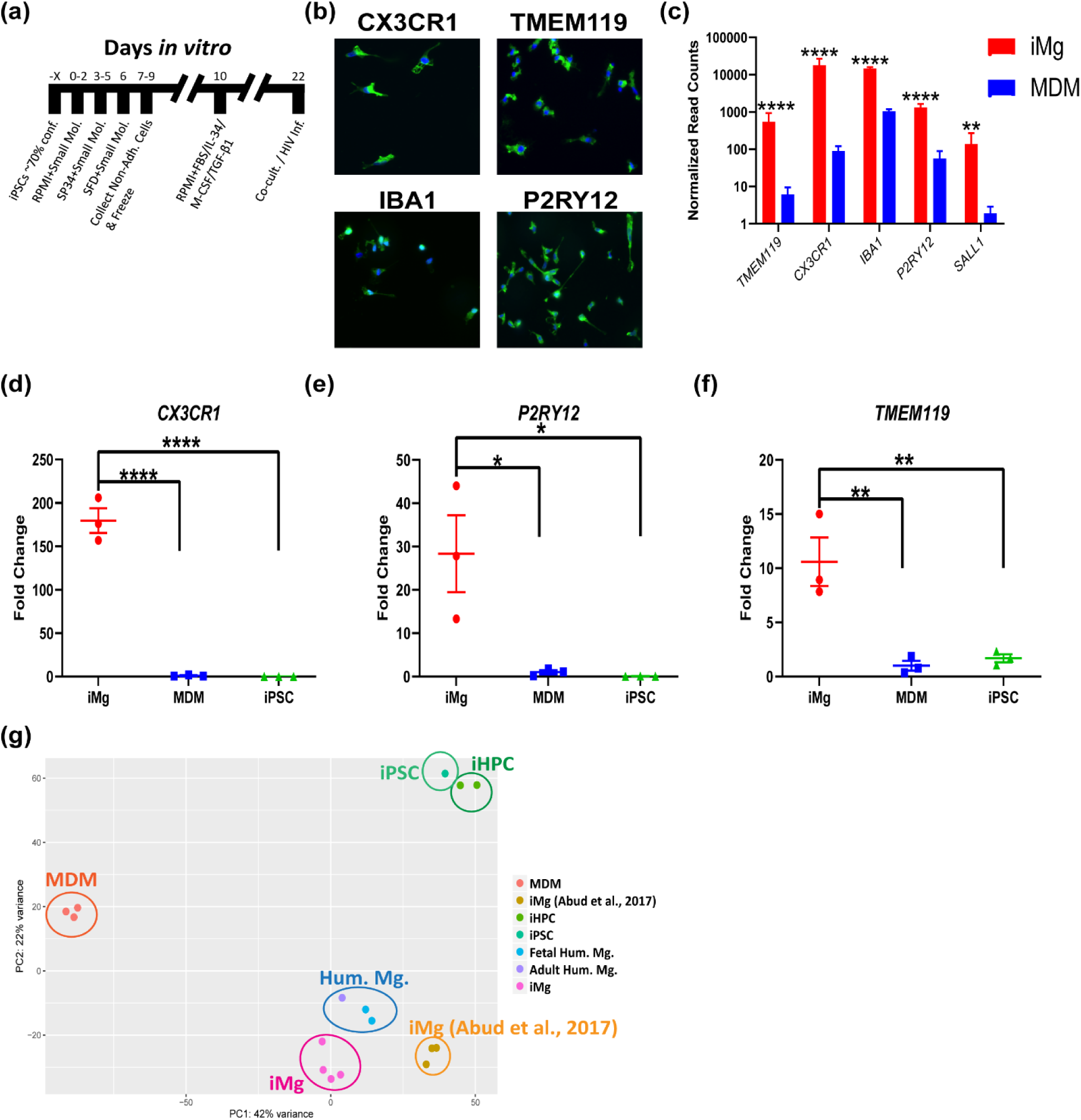
Generation and characterization of iMg. (a) Timeline for iMg differentiation and HIV infection. (b) Immunostaining for microglia specific markers: CX3CR1, TMEM119, IBA1, and P2RY12. All cultures were stained for DAPI. (c) Bulk RNAseq normalized read counts of specific microglia markers from iMg and MDMs. n=4 iMg, n=3 MDMs. Benjamini-Hochberg, FDR 0.01, **p<0.01, ****p<0.0001, error bars represent SEM. (d-f) qRT-PCR validation for expression of *CX3CR1* (d), *P2RY12* (e), and *TMEM119* (f) in iMg. Probes normalized to *GAPDH* expression. n=3 cell lines, one-way ANOVA, Dunnett’s post hoc analysis, *p<0.05; **p<0.01, ****p<0.0001, error bars represent SEM. (g) PCA for monocyte-derived macrophages, iMicroglia, and (Abud et al., 2017) iMg, iPSC, iHPC, and primary human microglia. Each dot represents a bulk RNAseq sample. MDM - monocyte-derived macrophage; iHPC – induced hematopoietic progenitor cells. iPSC, iHPC, Hum. Mg., iMg (Abud et al., 2017) samples were all retrieved from (Abud et al., 2017).

### iMg Are productively infected with HIV and respond to multiple antiretrovirals

We investigated the effects of HIV infection and ART in an iMg mono-culture. We first determined infectivity of the iMg. We exposed the iMg to 50ng/mL of the CSF-derived, R5-tropic JAGO strain of HIV (Chen et al., 2002). We performed a 15-day infection, based on prior studies of MDMs (O’Donnell et al., 2006) which resulted in 94.5 ± 5.5% of the iMg positive for the HIV capsid protein P24 (Figures 2a and 2b) and exhibited vast syncytia (Figure 2-figure supplement 1b). 89.1 ± 3.8% of P24(+) iMg were multi-nucleated (Figure 2c), and 100% of P24(-) iMg were single-nucleated (Figure 2d). Reverse transcriptase activity showed peak viral production occurred near day 15 (Figure 2e). iPSCs did not become infected at this concentration (Figure 2-figure supplement 2b). We infected four separate iMg lines; however, one was heterozygous for the CCR5Δ32 mutation, leading to impaired infection (Liu et al., 1996) (Figure 2-figure supplement 2c and 2d). Therefore, we excluded it from all future analysis. To our knowledge, this this the first time iPSC-derived microglia have been infected with HIV.

**Figure 2:**
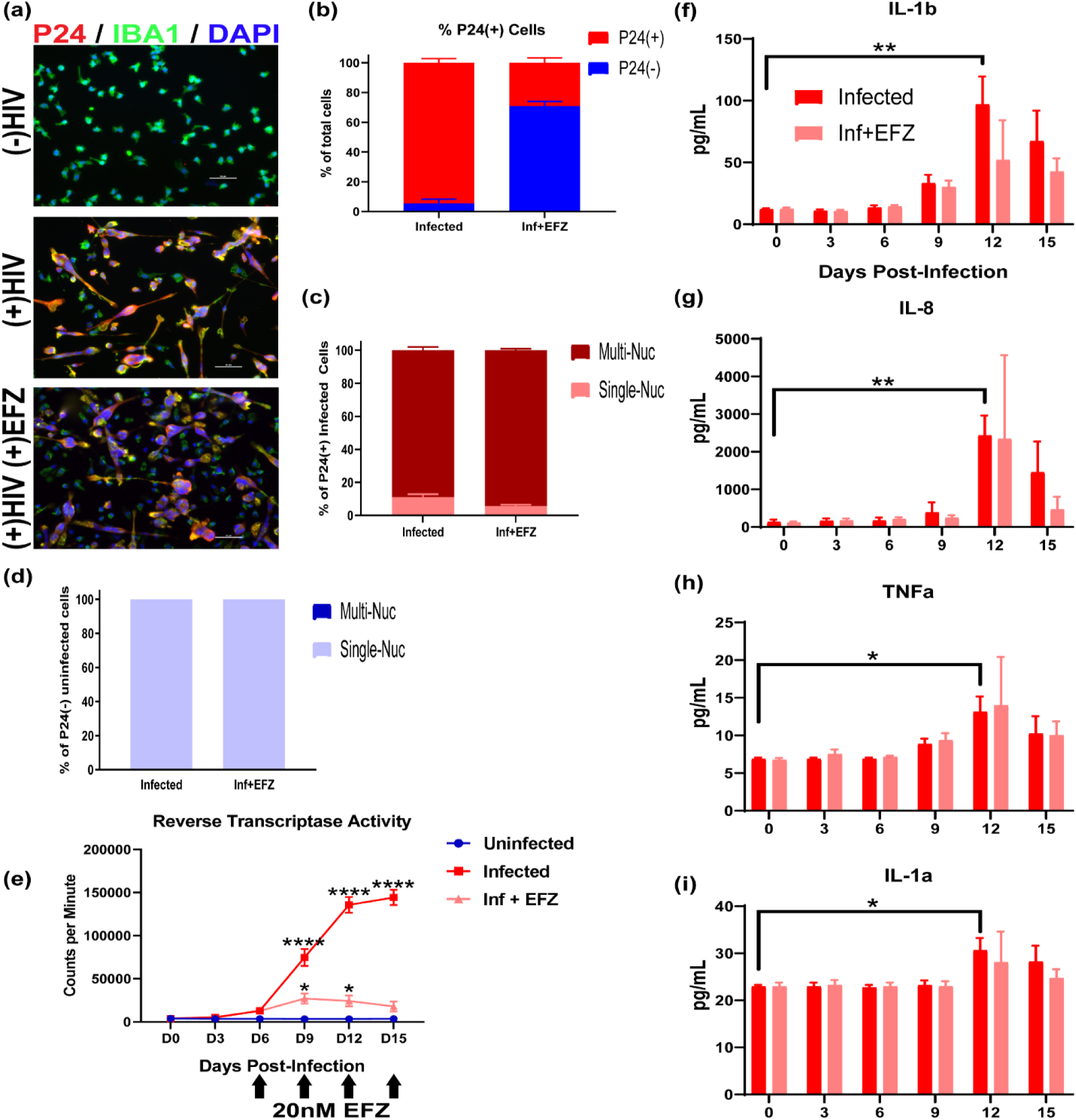
iMicroglia are infected by HIV, produce an inflammatory response, and respond to EFZ. (a) Immunostaining showing reduced percentage of P24+(red) IBA1+(green) iMg in HIV-infected mono-cultures + EFZ treatment compared to infected cultures with no EFZ treatment. (b) Percentage of P24+ cells in mono-culture iMg at D15 for infected and Inf+EFZ conditions. n=4 infections, error bars represent SEM. (c) Percent of P24+ single-nucleated and multi-nucleated iMg in mono-culture for infected and Inf+EFZ conditions. n=4 infections. (d) Percent of P24(-) single-nucleated and multi-nucleated iMg in mono-culture for Inf and Inf+EFZ conditions. n=4 infections. (e) Reverse transcriptase activity of Uninf, Inf, and Inf+EFZ (20nM) iMg show productive infection and response to EFZ. n=3 infections, one-way ANOVA, Dunnett’s post hoc analysis, *p<0.05; ****p<0.0001, error bars represent SEM. (f-i) Cytokine analysis of Infected iMg mono-culture displays increase in IL-1b (f), IL-8 (g), TNFa (h), and IL-1a (i) production in the infected iMg. n=4 infections, one-way ANOVA, Dunnett’s post hoc analysis, *p<0.05; **p<0.01, error bars represent SEM.

We tested two antiretroviral treatments that target different parts of the HIV life cycle, Darunavir (DRN) and efavirenz (EFZ) (Figure 2-figure supplement 2a). DRN is a protease inhibitor that prevents assembly of new virus (Wood & Flexner, 1998), and EFZ is a non-nucleoside reverse transcriptase inhibitor that prevents reverse transcription of the viral genome (De Clercq, 2004). We opted to focus on EFZ, as it is still a first line drug in many parts of the world (Best & Goicoechea, 2008; Taramasso et al., 2018), Further, because it blocks HIV infection at the reverse transcription phase, we can study non-productively infected, HIV-exposed cells alongside productively infected cells.. Importantly, the infected iMg responded as expected to EFZ (Figures 2a, 2b, and Figure 2-figure supplement 2a). At day 15, 29.2 ± 6.3% of the iMg were infected (Figures 2d and Figure 2-figure supplement 1c). 94.4 ± 1.72% of P24(+) iMg in the infected + EFZ mono-culture were multi-nucleated (Figure 2d), and 100% of P24(-) iMg in the infected + EFZ mono-culture were single-nucleated, same as the infected condition (Figure 2e).

### Infected iMg produced proinflammatory cytokines at peak infection, and EFZ treatment reduced the response

We also explored the changes in cytokine production over the course of infection ± EFZ. Infection led to increased production of several pro-inflammatory cytokines, specifically IL-1b, IL-1a, TNFa, and most prominently, IL-8 (Figures 2f-2i). However, the two other cytokines tested: IL-6 and IL-10, did not change across any condition (Figure 2-figure supplement 3a-3c). IL-8 production was increased in cultured, infected, fetal microglia (Tatro, Soontornniyomkij, Letendre, & Achim, 2014), and IL-1b RNA was increased in laser-captured microglia from brain samples of HIV-infected patients (Ginsberg et al., 2018). Infected iMg cultures exposed to EFZ had mitigated, non-significant responses to IL-1b, IL-8, TNFa, and IL-1a (Figures 2f-2i), suggesting a reduced inflammatory reaction. Uninf, Uninf+EFZ and the DMSO vehicle control did not elucidate a cytokine reaction across the six cytokines tested (Figure 2-figure supplement 3a-3e and Figure 2-figure supplement 4a-4d).

### Infected iMg have impaired cell cycle regulation and DNA repair and increased expression of inflammatory genes

Bulk RNAseq of three iMg lines ± HIV infection revealed significant changes in cell cycle regulation and DNA repair (Figure 2-figure supplement 5a). Infected immune cells will die quickly after infection (Cummins & Badley, 2010), and we observed a high percentage of giant multi-nucleated cells. Thus, it is unsurprising to see that most of the top canonical pathways altered involved cell cycle and DNA repair. While no inflammatory pathways were identified by IPA analysis, many inflammatory genes involved in the complement system, NF-κB signaling, and TNFa signaling were significantly upregulated (Figure 2-figure supplement 5c). In addition, other inflammatory genes including *IL1b*, *CCL8*, and *FOS* were also upregulated (Figure 2-figure supplement 5c).

Collectively, these data show that the iMg exhibit an inflammatory response similar to the that seen in the brain with increased production of IL-8, IL-1b, IL-1a, and TNFa. This inflammatory response is strongly attenuated with Efavirenz treatment. Bulk RNAseq analysis revealed changes in cell cycle and inflammation between uninfected and infected iMg, as expected.

### iNeurons were produced utilizing the NGN2 differentiation protocol

We utilized a published NGN2 protocol (Zhang et al., 2013) to generate a homogenous population of glutamatergic, forebrain-like excitatory neurons (Figure 3-figure supplement 1a and 1b). Symptoms of HAND include loss of working memory, fine motor skills, and executive function (Kathryn A. Lindl, Marks, Kolson, & Jordan-Sciutto, 2010; Saylor et al., 2016). Several of these functions are controlled by the prefrontal cortex and hippocampus, where most neurons are excitatory, glutamatergic neurons (Fuster, Bodner, & Kroger, 2000; Masterman & Cummings, 1997; Miller, 2000). We confirmed these neurons generate synapses (Figure 3-figure supplement 1c).

### iAstrocytes exhibit similar gene and protein expression as immature primary Astrocytes

We noticed that by day 2 of the NGN2 differentiation, the cells took on a neural precursor-like phenotype, expressing the neural progenitor markers Nestin and NCAM (Lendahl, Zimmerman, & McKay, 1990; Marmur et al., 1998), and the astrocyte marker SOX9 (Kang et al., 2012) (Figure 3-figure supplement 2a). Since neural progenitors can differentiate into astrocytes *in vivo*, we attempted to shift the neural differentiation at day 2 from neurons to astrocytes by removing the NGN2-inducing agent doxycycline, and adding FGF-2, EGF, and FBS to promote astrocyte differentiation (Figure 3a). After 90 days, a homogenous-appearing population of iAst were produced (Figure 3b). Bulk RNAseq revealed similar expression of several key genes for immature astrocytes between iAst and primary, fetal human astrocytes (Hu Ast) (Figure 3c and 3e). qRT-PCR validation of *THBS1* confirmed no difference in expression Hu Ast and iAst (Figure 1d). The overall gene expression profile of the iAstrocytes is similar to fetal human primary astrocytes (Figure 3e). We confirmed several of these genes at the protein level, including Nestin, Glutamine Synthetase, THBS1 and SOX9 (Figure 3b), with 84.5 ± 22.9% expressing SOX9, 97.4 ± 2.3% expressing Nestin, 91.3 ± 2.4% expressing Glutamine Synthetase, and 46.5 ± 24.8% expressing THBS1, an important protein in early synaptogenesis (Christopherson et al., 2005) (Figure 3b); however, they did not express GFAP (Figure 3-figure supplement 2b). Additionally, in mono-culture, they do not express the conventional glutamate transporter *GLT-1* or *GLAST* (Figure 3-figure supplement 2b and 2c). Although in tri-culture, scRNAseq showed expression of *GLT-1* (Figure 3-figure supplement 2e), which was not found in bulk RNAseq at the mono-culture stage. Neuronal activity regulates GLT-1 in astrocytes (Awabdh et al., 2016; Perego et al., 2000; Swanson et al., 1997), suggesting the iAst have a more *in vivo*-like phenotype when in the more physiologically relevant tri-culture.

**Figure 3:**
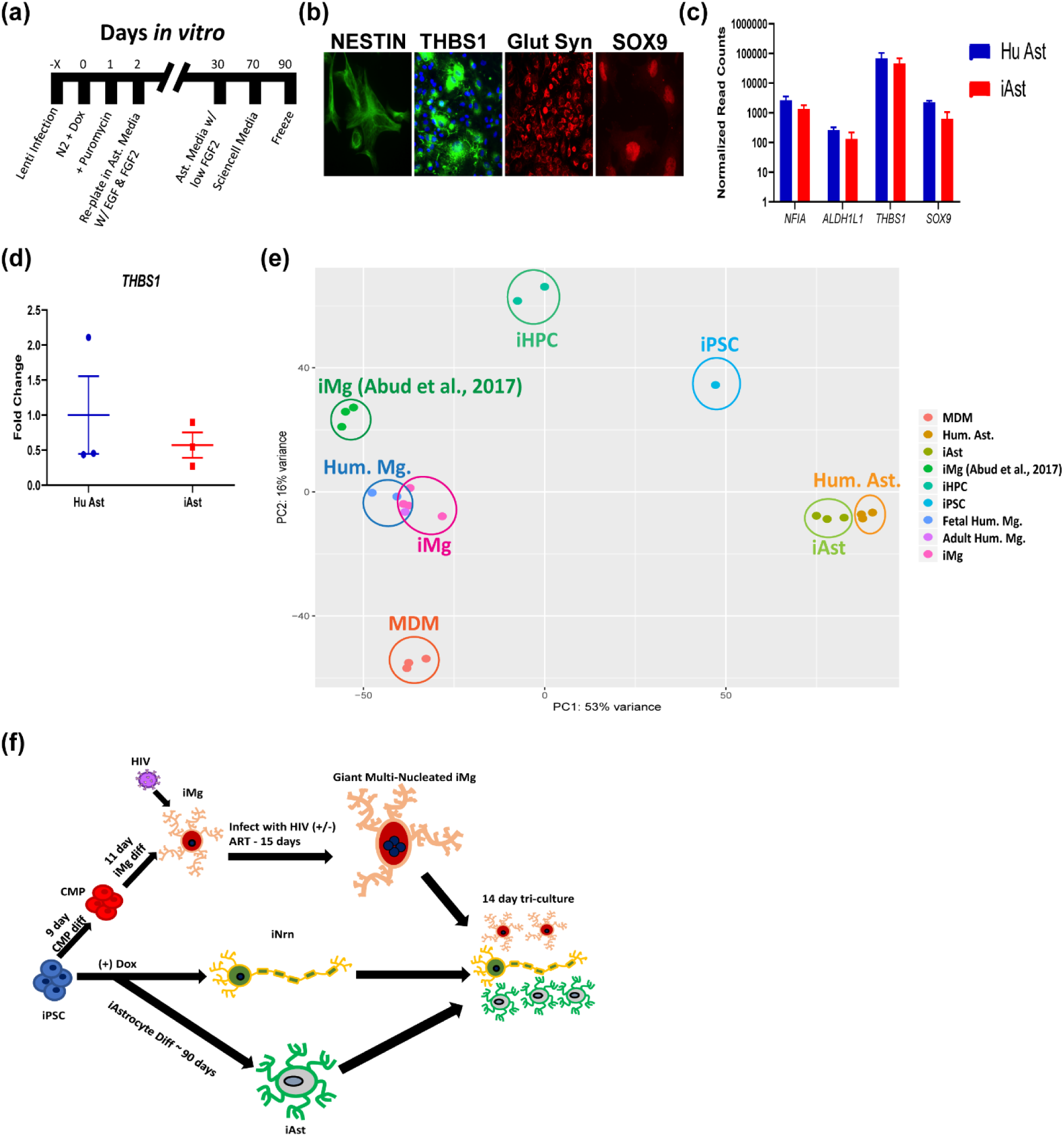
Generation and characterization of iAst and formation of tri-cultures. (a) Timeline for iAstrocyte differentiation. (b) Immunostaining for the Astrocyte specific markers: Nestin, Thrombospondin-1, Glutamine Synthetase, and SOX9. (c) Bulk RNAseq normalized read counts of select astrocyte specific genes from primary human astrocytes and iAst. n=3 cell lines, Benjamini-Hochberg, FDR 0.01, error bars represent SEM. (d) qRT-PCR validation for expression of *THBS1* in Hu Ast and iAst. Probe normalized to *GAPDH* expression. n=3 cell lines, two-tailed t test, error bars represent SEM. (e) PCA for monocyte-derived macrophages, iMicroglia, primary human astrocytes, and iAstrocytes, and (Abud et al., 2017) iMg, iPSC, iHPC, and primary human microglia. Each dot represents a bulk RNAseq sample. MDM - monocyte-derived macrophage; Hum. Ast. – primary fetal human astrocytes; iHPC – induced hematopoietic progenitor cells. iPSC, iHPC, Hum. Mg., iMg (Abud et al., 2017) samples were all retrieved from (Abud et al., 2017). (f) Flowchart for differentiations into iMg, iAst, and iNrn and combination into tri-culture ± HIV infection and ART.

### iAst respond to HIV-induced cytokines produced by the iMg

We also exposed the iAst in mono-culture to the most highly expressed cytokines in the HIV-infected iMg, to determine if the cytokines produced by infected iMg can elicit an inflammatory response in the iAst. We exposed the iAst to IL-1b and IL-8 at 10ng/mL for eight hours and then analyzed the supernatants on the same six-cytokine panel. We found that, of the six cytokines tested, the iAst produced increased amounts of IL-1a (Figure 2-figure supplement 5b), suggesting that the iMicroglia can elicit an inflammatory response in the iAst. In addition, it is interesting that IL-1a was released given its role in producing A1 astrocytes (Liddelow et al., 2017).

### The HiPSC-derived tri-culture is sustainable with iAst and iMg derived from novel differentiation protocols

In order to study the cell-cell interactions during HIV infection and subsequent antiretroviral treatment (ART), we developed a novel tri-culture system of HiPSC-derived microglia (iMg), neurons (iNrn), and astrocytes (iAst) by separately differentiating each cell type, infecting the iMg (± ART), then co-plating the different populations into a carpet culture for analysis. The tri-culture is able to survive for at least 2 weeks (Figure 3f).

### scRNAseq identified each of the three cell types in all four conditions

Next, we studied the cell types’ reactions in a more physiologically relevant tri-culture. To create the tri-culture, we began the iNrn differentiation, added the iAst at D5 of the iNrn differentiation, after the Ara-C had been removed, and added the iMg at D7 of the iNrn differentiation. The tri-culture was maintained for an additional 14 days, to D21 of the iNrn differentiation (Figure 4a). To assay gene expression changes in each of the three cell types during HIV infection ± EFZ, we conducted scRNAseq on the tri-culture under four conditions: Uninf, Inf, Inf+EFZ, and Uninf+EFZ (Figure 4a). Cells were sequenced from the Uninf (n=6,564), Inf (n=7,431), Inf+EFZ (n=7,111), and Uninf+ EFZ (n=10,071) conditions. All conditions were then aggregated and initially analyzed through the cellranger pipeline (10x Genomics, v.3.0.1). Secondary analysis was performed using the Seurat package in R. We generated 16 unbiased clusters (Figure 4b). First, we separated the clusters by cell type, then broke down each cell cluster by condition. We assigned clusters to one of the three cell types by expression of several key genes: iMg by expression of *IBA1*, *PU.1*, and *CD4*; iNrn by expression of *MAP2* and *SYN1*; and iAst by expression of *THBS1* and *SOX9* (Figures 4c-4e). There were four clusters (cluster 5, 10, 12, and 13) that did not fit into any of the three cell types by expression of the chosen markers (Figure 4b and 4f). To determine what the fourth cell type might represent, we examined the expression of the top 20 genes in each of the 4 cell types (Figure 4g). The gene expression pattern of the undesignated cells best matched the iAst, but scRNAseq did not capture expression of THBS1 or SOX9, suggesting these cells represent less mature versions of the iAst. Hence, they were excluded from further analysis (Figure 5a). We then separated each cell type by condition (Figure 5b).

**Figure 4:**
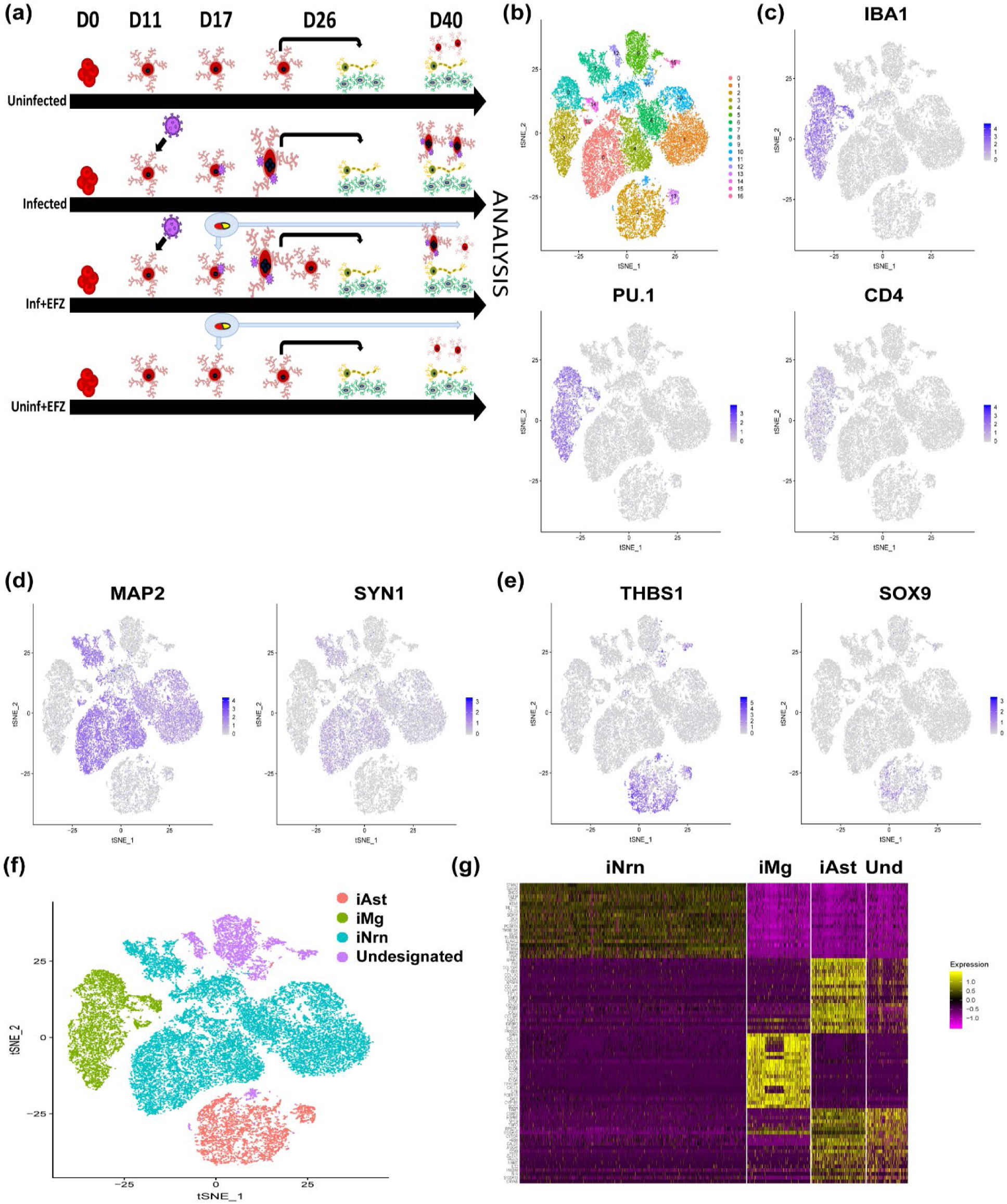
scRNAseq identified each of the three cell types in all four conditions. (a) Timeline from start of CMP differentiation through tri-culture for the four conditions. (b) t-SNE of unbiased clustering of combined scRNAseq from all four conditions. (c-e) expression patterns of cell type specific markers for microglia (c), neurons (d), and astrocytes (e). (f) t-SNE clustering by cell type based on cell type specific marker expression. One cluster did not align with any of the three cell types by the expression patterns chosen in (c-e). (g) Heat map of top 20 genes expressed in each cell type reveals the undesignated cluster of cells to be similar to iAst expression profile, suggesting they may be immature iAst.

**Figure 5:**
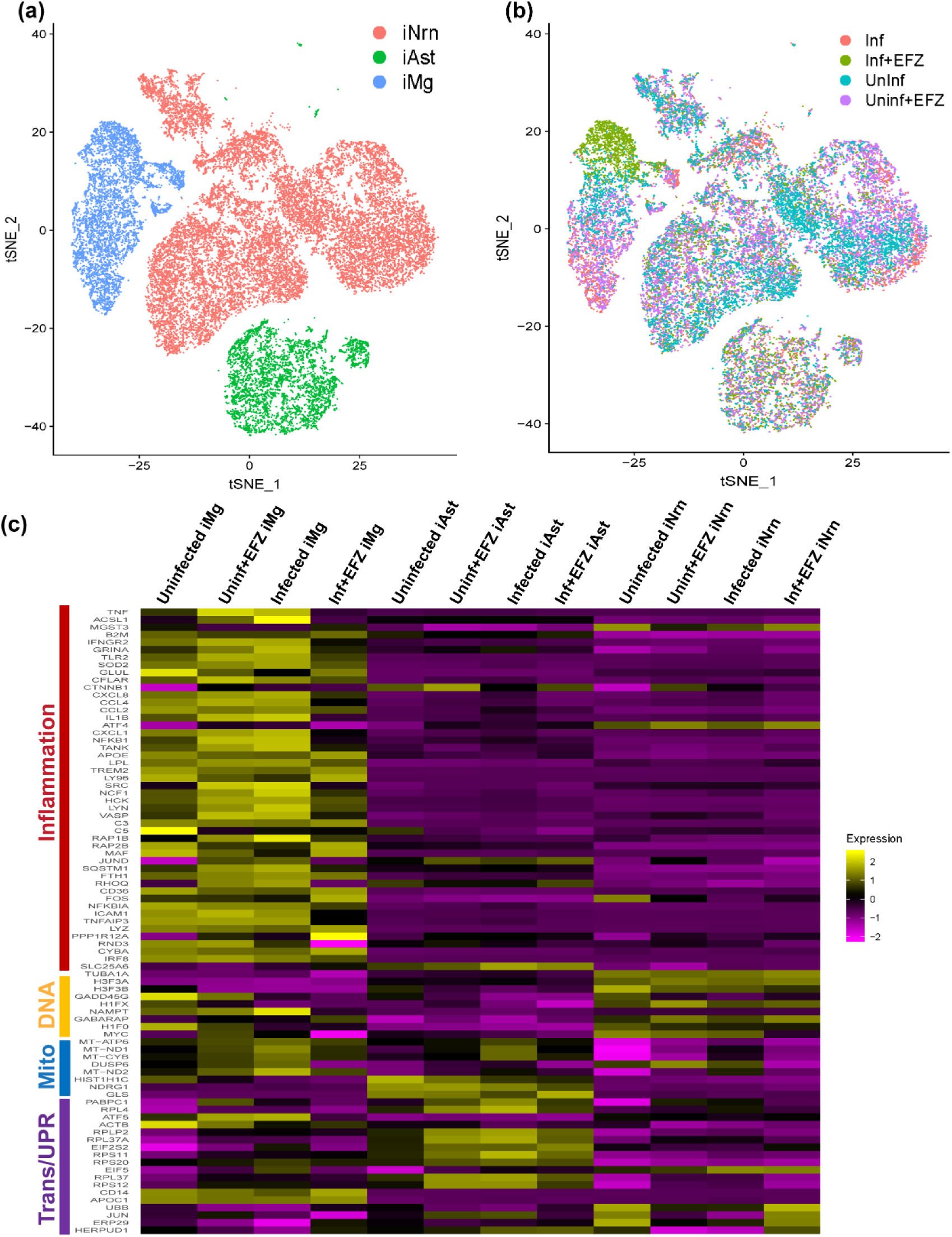
iMg exhibit the largest inflammatory response by gene expression among the three cell types in tri-culture. (a) t-SNE plot of all cells from all conditions excluding the undesignated cluster. (b) t-SNE from (a) broken down by condition. (c) Heatmap of inflammation, DNA accessibility, mitochondria, and translation/UPR related genes in iAst, iNrn, and iMg in all four conditions: Uninf, Uninf+EFZ, Inf, Inf+EFZ.

### iMg exhibit the largest inflammatory response by gene expression

After the scRNAseq was separated by cell type (Figure 5a) and condition (Figure 5b), We examined a panel of inflammation, DNA accessibility, mitochondria, and translation/UPR-related genes to determine which of the three cell types had the greatest changes in relevant gene expression (Figure 5c). The iAst had the most prominent change in translation and UPR related genes, with increased expression of several ribosomal genes and *EIF2S2*, a subunit of EIF2, in the Inf condition compared to Uninf. Whereas, the iNrn had the largest increases in DNA accessibility genes, including several histone-related genes. Not surprisingly, the iMg population showed the highest expression of inflammatory and mitochondria related genes. Inf iMg had increased expression of several inflammatory cytokine genes, including *CXCL8* and *IL-1b*. This corroborates with the increased production if IL-8 and IL-1b in infected iMg in mono-culture. The Inf+EFZ iMg had lower expression levels than Inf iMg for these genes, again corroborating with the mono-culture cytokine production. Although, the Inf+EFZ iMg had increased expression of the inflammatory genes *FOS* and *APOE*, compared to control. These results suggest that the iMg are mostly responsible for the inflammatory response during infection ± EFZ; however, there are distinct responses ± EFZ. To further investigate the differences, we next performed pairwise comparisons of each cell type across all four conditions.

### Inflammatory pathways and EIF2 signaling were consistently dysregulated among all three cell types during infection, but iMg were the most affected

We first compared the Inf condition to Uninf. Several inflammatory pathways were highly activated in Inf iMg compared to Uninf iMg, including Il-8 and NF-κB signaling. One of the top pathways dysregulated in Inf iMg compared to Uninf was the EIF2 pathway (Figures 6b). EIF2 signaling is involved in the UPR and, more broadly, the ISR (Cui, Li, Ron, & Sha, 2011; Janssens, Pulendran, & Lambrecht, 2014; Walter & Ron, 2011). The EIF2 pathway was not only dysregulated in iMg, but also was consistently increased in the iAst and iNrn (Figure 6-figure supplement 1a and 1b). However, only iMg had increased expression of *ATF4*, the downstream transcription factor of the PERK arm of the ISR, which leads to increased cytokine production (Masuda, Miyazaki-Anzai, Levi, Ting, & Miyazaki, 2013) in Uninf v Inf (Figure 5c). Previously, we have shown that the PERK arm of the ISR is activated in neurons and astrocytes from human brain samples with HAND (C. Akay et al., 2012; K. A. Lindl, Akay, Wang, White, & Jordan-Sciutto, 2007).

**Figure 6:**
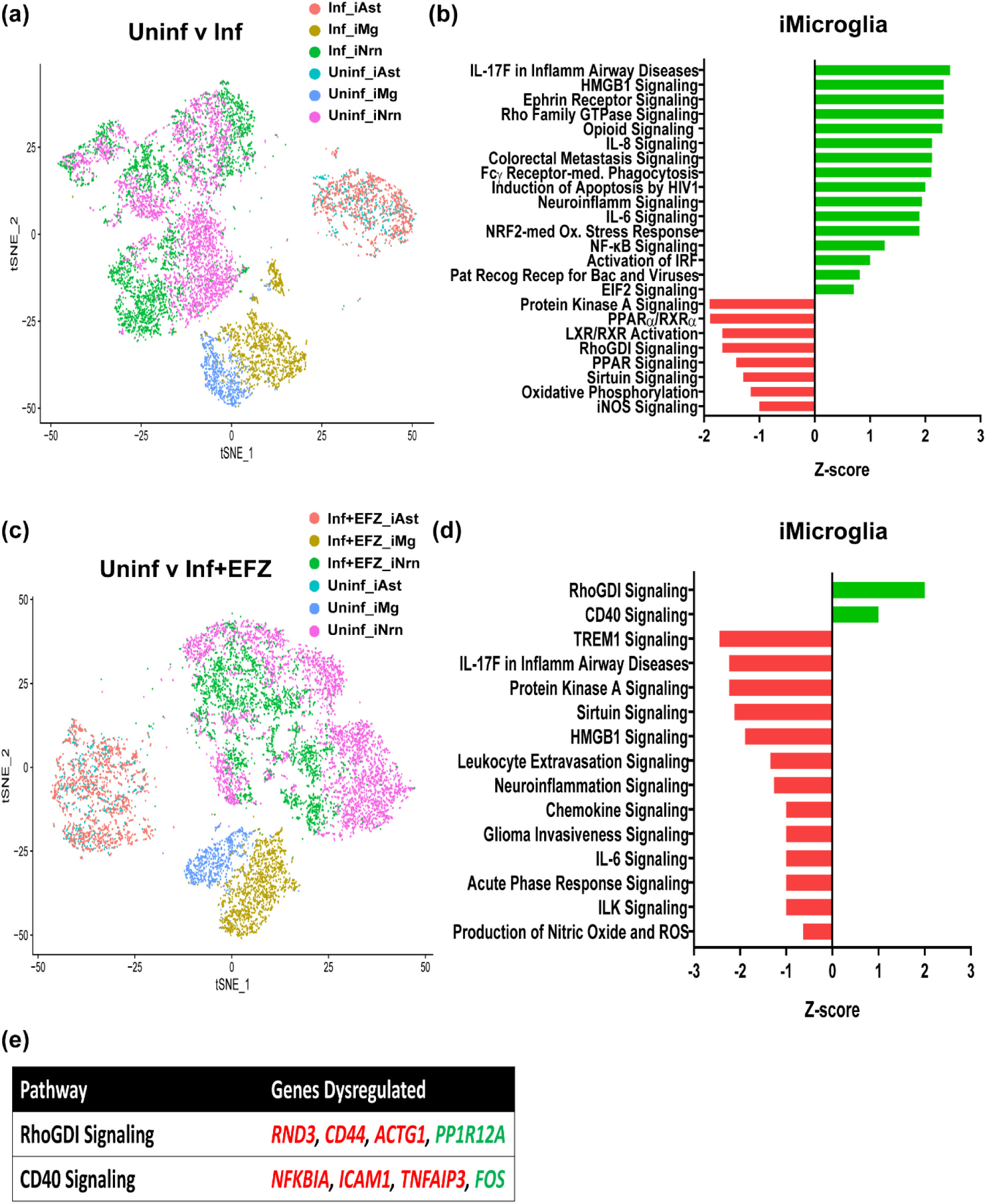
iMg activate RhoGDI and CD40 in response to HIV infection with EFZ treatment. (a) t-SNE plot of Uninf and Inf conditions. (b) Ingenuity pathway analysis of iMg between Uninf and Inf conditions. Uninf is baseline. Benjamini-Hochberg FDR 0.05, Fisher’s exact <0.05, z-score cutoff ± 0.5. (c) t-SNE plot of Uninf and Inf+EFZ conditions. (d) Ingenuity pathway analysis of iMg between Uninf and Inf+EFZ conditions. Uninf is baseline. Benjamini-Hochberg FDR 0.05, Fisher’s exact <0.05, z-score cutoff ± 0.5. (e) Specific genes dysregulated that are involved in the RhoGDI and CD40 pathways in Inf+EFZ iMg compared to Uninf iMg. Red genes are downregulated. Green genes are upregulated.

### Infection+EFZ had distinct increased activation of RhoGDI and CD40 pathways

We next compared Uninf to Inf+EFZ, expecting to see a dampened immune response compared to Inf. Many of the top affected pathways in Inf iMg were related to inflammation; however, there does seem to be a stark difference between Inf iMg and Inf+EFZ iMg (Figures 6b and 6d). The Inf+EFZ had a much milder inflammatory reaction, where RhoGDI and CD40 signaling were the only upregulated pathways compared to Uninf iMg (Figure 6d and 6e). RhoGDI negatively regulates Rac, which functions in multiple inflammation pathways (Wilkinson & Landreth, 2006), and CD40 activates NF-κB signaling (D’Aversa et al., 2008; Homig-Holzel et al., 2008). A milder inflammatory response corroborates well with lesser disease severity seen in HAND patients who are taking ART (Saylor et al., 2016).

The scRNAseq data suggest the microglia were most affected by the infection ± EFZ, with major changes to EIF2 signaling, inflammatory, oxidative damage, and phagocytic gene pathways. However, the distinct activation of RhoGDI and CD40 in the Inf+EFZ condition suggests the combination of Inf+EFZ creates a distinct effect. We thus further investigated the effects of Inf+EFZ treatment compared to Inf and Uninf+EFZ treatment.

### Inf+EFZ has an attenuated, but distinct inflammatory response compared to Inf and Uninf+EFZ

Inf iMg had increased activity in several inflammatory pathways including IL-6 signaling, neuroinflammation signaling, and Fcγ receptor-mediated phagocytosis compared to Inf+EFZ (Figure 6-figure supplement 2a and 2b), suggesting the Inf+EFZ had an overall lower immune response. However, the EIF2 signaling pathway, part of the UPR, was lower in Inf iMg compared to Inf+EFZ (Figure 6-figure supplement 2b).

We next compared Uninf+EFZ to Inf+EFZ (Figure 6-figure supplement 2c). EIF2 signaling was increased in the Inf+EFZ iMg (Figure 6-figure supplement 2d), similar to the Inf and Inf+EFZ comparison (Figure 6-figure supplement 2b). However, inflammatory pathways including neuroinflammation and ROS/iNOS signaling were higher in Uninf+EFZ compared to Inf+EFZ iMg (Figure 6-figure supplement 2d). Inf+EFZ iMg again had increased activation of the CD40 and RhoGDI pathways (Figure 6-figure supplement 2d and 2e) compared to Uninf+EFZ, suggesting a distinct inflammatory response with the combination of infection and EFZ treatment.

iAst had activation of the neuroinflammation and RhoGDI signaling pathways in Inf+EFZ than Uninf+EFZ (Figure 7-figure supplement 1e). These results show a stark difference in responses to EFZ treatment alone and EFZ treatment with infection, suggesting combinatorial and probably interacting effects of infection and EFZ treatment.

### Uninf+EFZ creates an inflammatory response similar to Inf

Inf+EFZ had a starkly different response than Uninf+EFZ. So, we further investigated the effect of EFZ treatment alone compared to Uninf and Inf treatments. ART can be used as a preemptive treatment against HIV infection; however, it has various inflammatory and neurotoxic side effects (Cagla Akay et al., 2014; Robertson, Liner, & Meeker, 2012). How ART affects different cell types in co-culture has not been explored. We compared Uninf+EFZ to the Uninf (Figures 7a, 7b, and Figure 7-figure supplement 1a and 1b) and Inf (Figures 7c, 7d, Figure 7-figure supplement 1c and 1d) and found a surprisingly similar gene expression profile to the Inf condition, especially in the iMg and iAst. The pairwise comparison of Uninf and Uninf+EFZ revealed multiple inflammatory pathways increased in Uninf+EFZ iMg, with neuroinflammation signaling as the most activated pathway (Figure 7b). Several of the other pathways activated, including IL-6 and IL-8 signaling, were also found in the Uninf v Inf comparison (Figure 6b). EIF2 signaling was increased in Uninf+EFZ iNrn compared to Uninf (Figure 7-figure supplement 1b), similar to Inf iNrn (Figure 6-figure supplement 1b). EIF2 signaling was not increased in the iAst, unlike in the Inf comparison, and there was a reversal for sirtuin signaling with inhibition in the Uninf+EFZ versus Uninf (Figure 7-figure supplement 1a), suggesting Uninf+EFZ elicits a similar, but not identical inflammatory response as Inf.

**Figure 7:**
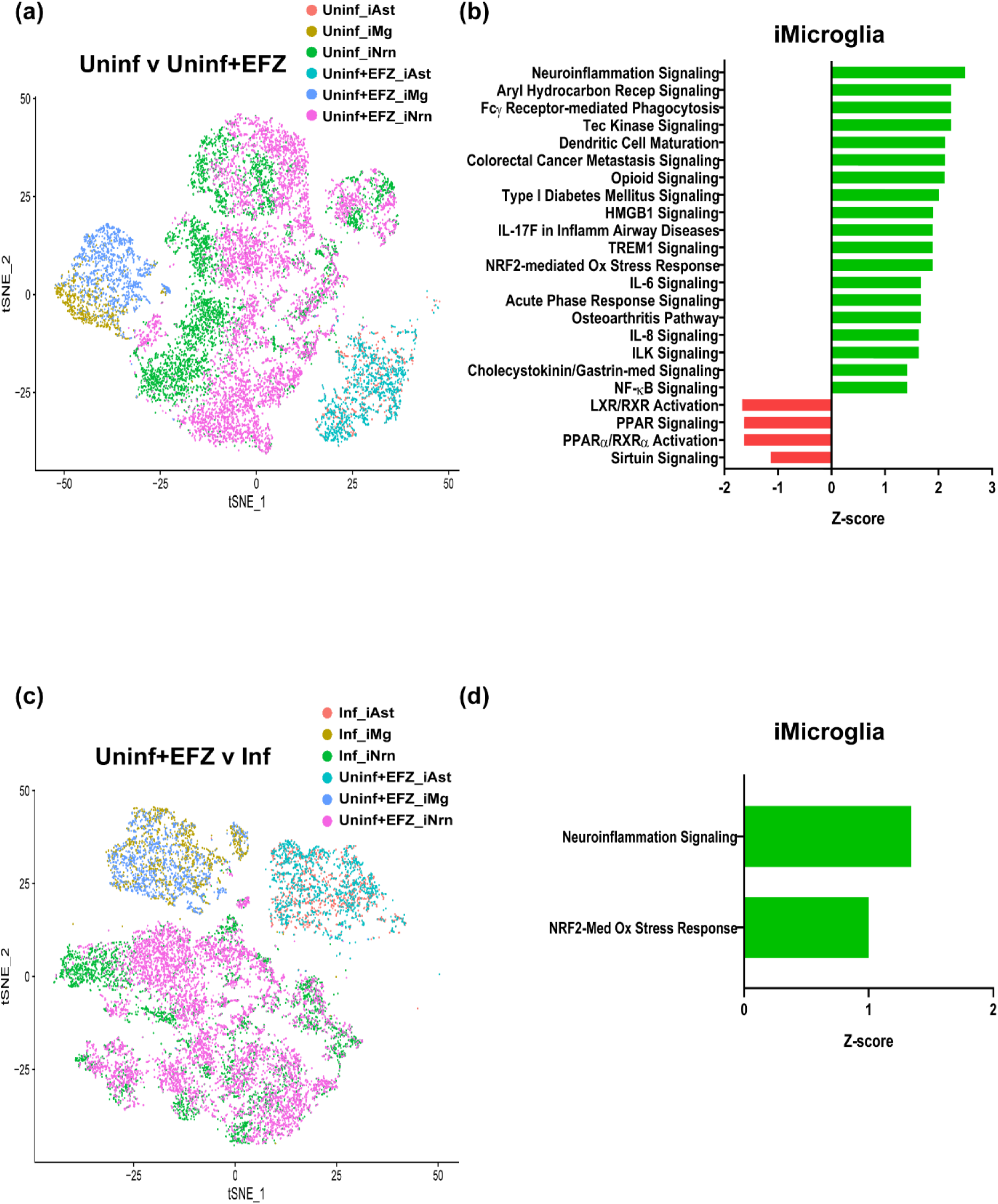
EFZ treatment alone elicits comparable response as infection. (a) t-SNE plot of Uninf and Uninf+EFZ conditions. (b) Ingenuity pathway analysis of iMg between Uninf and Uninf+EFZ conditions. Uninf is baseline. Benjamini-Hochberg FDR 0.05, Fisher’s exact <0.05, z-score cutoff ± 0.5. (c) t-SNE plot of Uninf+EFZ and Inf conditions. (d) Ingenuity pathway analysis of iMg between Uninf+EFZ and Inf conditions. Uninf+EFZ is baseline. Benjamini-Hochberg FDR 0.05, Fisher’s exact <0.05, z-score cutoff ± 0.5.

We found iMg had higher activation of neuroinflammation signaling and NRF2-mediated oxidative stress response in Inf compared to Uninf+EFZ (Figures 7c and 7d). iNeurons had higher oxidative phosphorylation activation; however, we found no statistically significant changes in any canonical pathways in iAst between the conditions (Figure 7-figure supplement 1c and 1d), suggesting an inflammatory response by iMg and change in mitochondria function in neurons in response to HIV infection of the former.

Infection, EFZ treatment alone, and infection with EFZ treatment all induced distinct responses not only across conditions, but also across cell types within conditions. These data reveal a potential downside to pre-emptive ART use and suggest that the inflammatory response from Infected to Inf+EFZ to Uninf+EFZ is not a linear regression as one might expect, but rather three distinct responses. We further investigated these gene pathway changes by testing for cytokine production and phagocytic abilities.

### Synaptophagocytosis is impaired in Inf, Inf+EFZ iMg, and Uninf+EFZ iMg

We interrogated the ability of iMg to phagocytose synaptic material to further explore the possible differences in the responses to infection ± EFZ and EFZ treatment alone. Fcγ receptor mediated phagocytosis pathway was activated in the Inf iMg as well as the Uninf+EFZ iMg compared to Uninf iMg (Figures 6b and 7b). Aberrant synaptophagy by microglia has been implicated in multiple neurological disorders including Alzheimer’s and Frontotemporal dementia (Hong et al., 2016; Lui et al., 2016). Many antiretrovirals, including Efavirenz, have reported neurotoxic effects (Ciccarelli et al., 2011; Robertson et al., 2012) and reduce phagocytosis (Giunta et al., 2011). In addition, HIV-infected macrophages and uninfected macrophages exposed to HIV display decreased phagocytic capabilities, specifically through the Fcγ receptor-mediated pathway and not complement, due to Nef and Tat inhibiting endocytosis (Debaisieux et al., 2015; Giunta et al., 2008; Mazzolini et al., 2010), but human microglia synaptophagocytosis has never been tested.

In order to determine their ability to phagocytose synapses, we generated tri-culture of the iMg with the iNrn and iAst and examined them 14 days after adding the iMg (Figure 4a). The cultures were subjected to immunofluorescence for IBA1, the lysosomal marker LAMP1, and the pre-synaptic marker SYN1. We found colocalization of SYN1 in the LAMP1+ lysosomes of the iMg (Figure 8a and 8b), suggesting that the iMg phagocytize synaptic material. We confirmed that iMg are in contact with iNrns (Figure 8c), and importantly, only the iMg are infected as only they, and not the iNrns (Figure 8d) or iAstrocytes, show P24 immunofluorescence. Additionally, RT activity confirmed that the iMg in the infected condition remain productively infected (Figure 9a). We were also able to delineate infected from uninfected iMg in the Inf+EFZ condition by multi-nucleation (Figures 2c, 2d, and Figure 8-figure supplement 1a).

**Figure 8:**
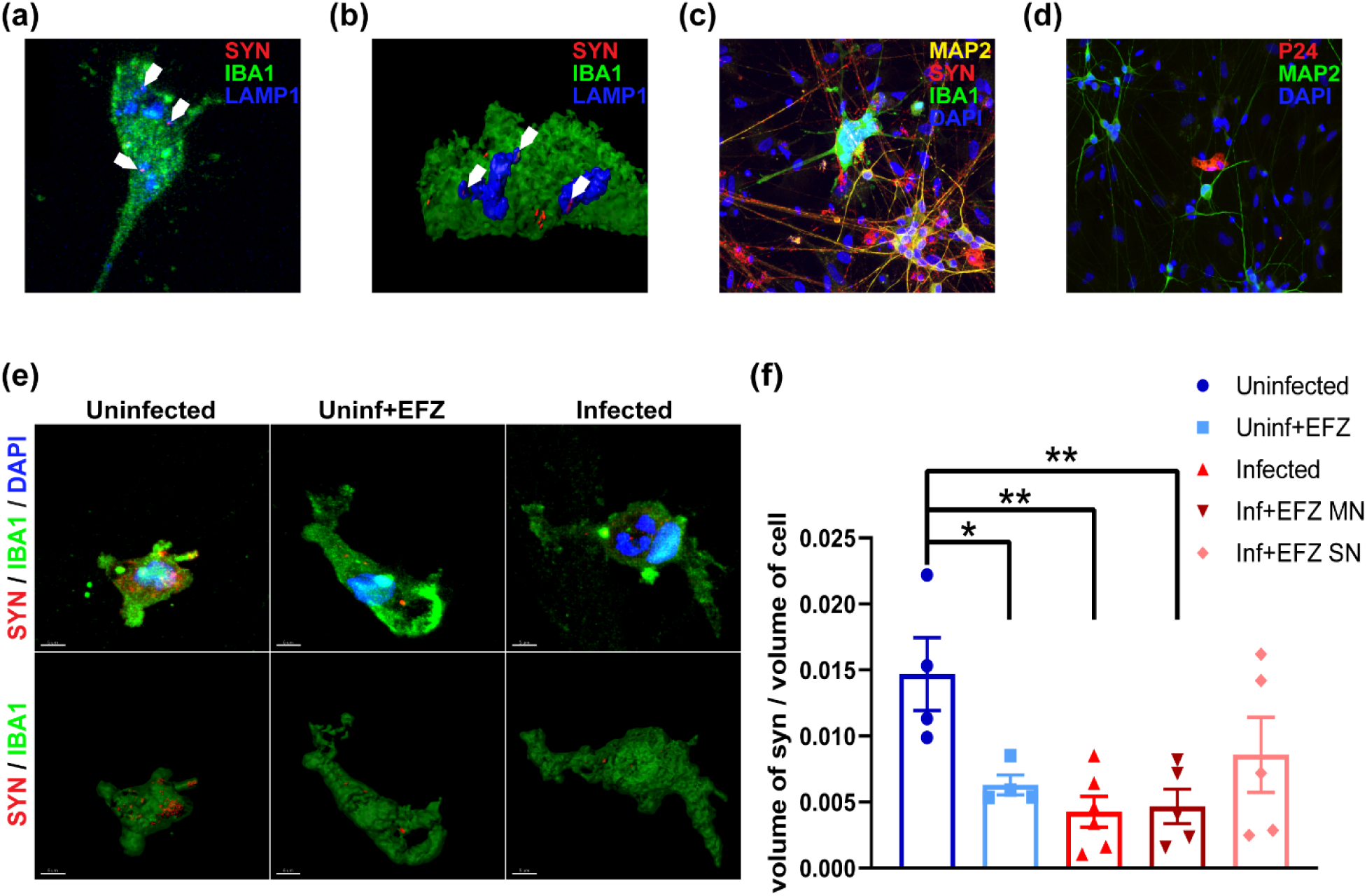
HIV-infected iMicroglia have reduced synaptophagocytosis, which is ameliorated by EFZ treatment. (a&b) Immunostaining (a) and surface reconstruction of a side view (b) of iMg in an iNrn co-culture displaying synaptic phagocytosis by co-localization of synaptophysin+ (red) puncta within LAMP1+ (blue) lysosomes in IBA1+ (green) iMg. (c) Giant multi-nucleated IBA1+ (green) iMg potentially interacting with synaptophysin+ (red) synapses on MAP2+ (yellow) dendrites. (d) Multi-nucleated iMg, but not MAP2+ (green) iNrn or iAst, are P24+ (red) in the tri-culture after 14 days. (e) Representative surface reconstructions of synaptophysin signal (red) within IBA1+ (green) iMg in the Uninf, Uninf+EFZ, and Inf tri-cultures. SN-single nucleated. MN-multinucleated. Scale bar represents 5µm. (f) Synaptophagy is significantly decreased in infected iMg ± EFZ and uninfected iMg + EFZ compared to control. n=4 (4 Uninf, 4 Uninf+EFZ, 6 Inf, 5 Inf+EFZ) differentiations, one-way ANOVA, Dunnett’s post hoc analysis, *p<0.05, error bars represent SEM.

**Figure 9:**
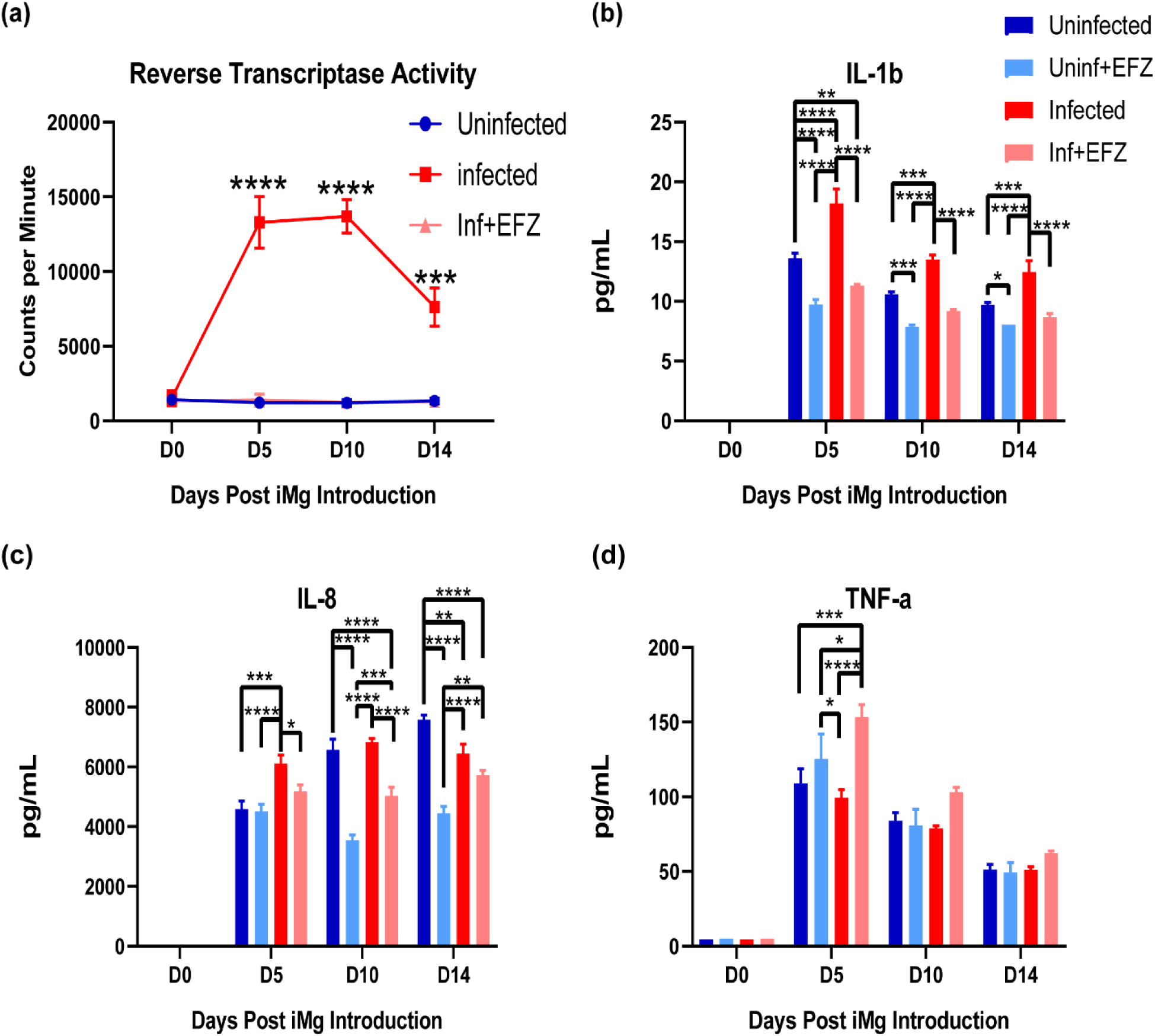
Inf+EFZ mitigates IL-1b and IL-8 production, but increases TNFa production. (a) Reverse transcriptase activity of Uninf, Inf, and Inf+EFZ tri-culture shows continual productive infection. n=3 infections, one-way ANOVA, Dunnett’s post hoc analysis, ***p<0.001; ****p<0.0001, error bars represent SEM. (b-d) Cytokine analysis of Uninf, Uninf+EFZ, Inf, and Inf+EFZ iMg displays increase in IL-1b (b) and IL-8 (c) production in the Inf tri-culture, and Inf+EFZ had increased TNFa (d). n=3 infections, two-way ANOVA, Tukey’s post hoc analysis, *p<0.05; **p<0.01; ***p<0.001; ****p<0.0001, error bars represent SEM.

We utilized confocal imaging and 3D reconstruction to measure the total synaptic volume, as indicated by synaptophysin immunofluorescence, within the iMg. The Inf iMg showed reduced synaptophagocytosis compared to Uninf iMg. In addition, in the Inf+EFZ condition, the infected (multinucleated) have reduced phagocytosis; however, the uninfected (single-nucleated) iMg in the Inf+EFZ condition trended lower but was not statistically significant. Lastly, Uninf+EFZ condition iMg also had impaired synaptophagocytosis (Figure 8e and 8f). Multiple factors from the virus itself, the immune response, and the effects of antiretrovirals are acting on the iMg during HIV infection. All these factors may play a role in reducing synaptic phagocytosis and warrant future study.

While the Fcγ receptor-mediated phagocytosis pathway was activated in Inf iMg, several of the genes dysregulated including: *ACTB*, *ACTR3*, *ARPC2*, *FYB1*, *RAC1*, and *VASP* (data not shown), were associated with the nascent phagosome and phagosome formation, which is targeted by Nef in impairing phagocytosis (Dirk et al., 2016; Mazzolini et al., 2010), suggesting a compensatory increase in gene expression for genes associated with the target for Nef, early endocytosis. This finding may partially explain the apparent paradox in increased pathway activity of Fcγ receptor-mediated phagocytosis, but impaired phagocytosis in Inf iMg.

To confirm that differences in apparent iMg synaptophagy are not secondary to reduced numbers of synapses overall in a given culture condition, we also measured local and random (50 µm radius areas with no iMg) synapse density to ensure the uninfected microglia were not in a more synapse dense area (Figure 8-figure supplement 2a and 2b). In fact, there was no significant difference in iMg-proximal or random synapse density across all conditions (Figure 8-figure supplement 2c and 2d). This finding aligns with previous studies that demonstrate inhibition of phagocytosis by viral proteins Tat and Nef in both infected and uninfected macrophages, (Debaisieux et al., 2015; Giunta et al., 2008; Mazzolini et al., 2010), and is also consistent with ART-related impairment of phagocytosis (Giunta et al., 2011).

### EFZ reduces the production of IL-8 and IL-1b by infected iMg, but enhances TNFa

Lastly, we compared cytokine production between each condition in tri-culture. Starting at the time of iMg addition, we collected supernatant every 5 days until day 14 of tri-culture. IL-8 and IL-1b were significantly increased in the infected tri-culture compared to uninfected control. IL-1b remained increased over uninfected; however, IL-8 decreases by day 10 (Figure 9b and 9c). Unlike in mono-culture, the Inf+EFZ condition had reduced cytokine production of IL-8 and IL-1b compared to uninfected. Uninf+EFZ had the lowest levels of cytokine production across all the groups (Figure 9b-9d). In addition, there was no increase in TNFa in the Inf tri-culture; however, the Inf+EFZ condition did have increased TNFa (Figure 9d). The Inf and Inf+EFZ tri-cultures also did not exhibit increases in IL-6 or IL-10, similar to our mono-culture findings (Figure 2-figure supplement 3c and 3d). These data combined with the scRNA data suggest there are distinct influences on the inflammatory response during infection alone and infection with EFZ. iMg in the Inf condition exclusively had increased expression of *IL-8* and *IL-1b* genes (Figure 5c). In addition, IL-8 and NF-κB signaling were activated only in iMg in the Inf condition by scRNAseq (Figure 6b) Interestingly, TNFa regulates *AP-1* (*FOS*) (Clark, Vagenas, Panesar, & Cope, 2005), a CD40 signaling pathway gene that is upregulated in Inf+EFZ iMg when compared to the Uninf (Figures 5c and 6d). These results further suggest distinct inflammatory reactions from Inf and Inf+EFZ that are mostly caused by the iMg.

## Discussion

We describe, to our knowledge, the first HiPSC tri-culture model to investigate the interdependent and individual roles of microglia, astrocytes, and neurons in the context of HIV infection. In recent years, there have been several new differentiation protocols for these cell types (Abud et al., 2017; Santos et al., 2017; Zhang et al., 2013) and several iPSC/primary cell co-cultures that have studied various neurological disorders (Lin et al., 2018; Park et al., 2018). This model goes one step further, exploring gene expression changes with scRNA sequencing in the context of viral infection and relevant drug treatments to begin to unfurl a particularly difficult disease to study.

Although there are missing components to this model, it is still able to recapitulate several findings *in vivo*, such as increases in IL-8 and IL-1β production and EIF2 signaling (C. Akay et al., 2012; Ginsberg et al., 2018; Tatro et al., 2014). It has also discovered novel changes that require further investigation but could lead to advancements in therapeutics. This system allowed us to identify sirtuin pathway dysregulation, which has been implicated in aging and Alzheimer’s, but not HAND (Cho et al., 2015; Jesko, Wencel, Strosznajder, & Strosznajder, 2017).

We were also able to show that the microglia seem to be the main culprit in the initial response to HIV infection; however, at least within the two-week period of this study, iMg infection does not lead to aberrant phagocytosis of synapses. This falls in line with previously reported impaired phagocytosis by HIV-infected or exposed macrophage (Debaisieux et al., 2015; Mazzolini et al., 2010). With Nef and Tat targeting early stages of the phagocytosis, and most of the genes dysregulated in the Fcγ receptor-mediated phagocytosis target early endocytosis, it is likely that the increased pathway activation is a compensation mechanism. Infection with HIVΔNef could confirm this.

In addition, we discovered a starkly different immune response at the gene and functional level in Inf versus Inf+EFZ. CD40 and RhoGDI activation could be controlled by the increased production of TNFa, that was only found in the Inf+EFZ condition. This persistent immune activation highlights the need for further studies into the role that antiretrovirals play in propagating the chronic inflammation seen in HIV patients today (Kolson, 2017; Saylor et al., 2016). The mitigated reaction we observed with EFZ treatment is consistent with human studies showing a reduced severity of HAND with the widespread use of antiretroviral therapies (Saylor et al., 2016), but also with its persistence despite control of viral load. Further, we revealed a substantial immune response to EFZ treatment alone. This warrants further study as people take ART as a preventative measure (Spinner et al., 2016).

HiPSC cultures are particularly useful for studying HIV neuropathology, since human primary neuronal cells and postmortem tissue are limited in availability and not amenable to molecular manipulation. In addition, HIV only infects human cells, rendering the interpretation of results from animal models more convoluted. Hence, mechanistic studies of the influences of HIV and ART on human neural cells are limited. This tri-culture system allows us to better study the mechanisms of early HIV infection in the brain; however, there are caveats to this system that must be considered. Each of the iPSC-derived cell types are similar to their *in vivo* counterparts by gene and protein expression, as well as, function, but are not exact. In addition, the iCells’ gene expression profile at the end stage of differentiation is relatively immature and more closely represents early stages of development *in vivo*. Our culture system allows reductionist study of three key cell types over weeks but may be refined by inclusion of additional cell types, and further optimization to model HAND in the adult setting.

This tri-culture has validated several findings in the field as well as produced multiple novel findings for HIV neuropathology. However, this model is not restricted to HIV neuropathology and can also be utilized to study other neurological disorders. This highly tractable, reductive system can be genetically and pharmacologically modified at any stage. These differentiations could also be used on patient-derived cells, creating a disease-relevant, patient-specific tri-culture system. Similar cultures have been developed (Haenseler et al., 2017; Park et al., 2018), but not to this complexity, as rapid a differentiation, or with a focus on viral infections. The tri-culture can be another, instrumental tool in understanding the workings of complex neurological disorders and in developing novel therapeutic strategies.

## Materials and Methods

### Human induced pluripotent stem cell lines

HiPSC lines for the iNrn and iAst were generously received from Herbert M. Lachman, MD., Einstein University, Bronx, New York. All lines were trained over to a feeder-free system with Stem MACS iPS-Brew XF media (Miltenyi Biotec 130-104-368). Lines were tested for mycoplasma using Lookout mycoplasma PCR detection kit (Sigma MP0035). iPSC lines for the iMg were cultured by the Human Pluripotent Stem Cell Core (CHOP).

### iNeuron differentiation of iPSCs

iPSCs were transfected with two plasmids VSVG.HIV.SIN.cPPT.CMV.mNgn2.WPRE and VSVG.HIV.SIN.cPPT.CMV.rTTA.WPRE, produced by Marius Wernig (Stanford University) and packaged by the University of Pennsylvania Viral Vector Core. Cells were exposed to 1ug/mL polybrene (Sigma Aldrich TR-1003). Media is fully exchanged six hours after exposure. iPSCs are differentiated according to a previously published method (Zhang et al., 2013). In brief, after transfection, iPSCs were exposed to N2 media containing 5 mL N2 Supplement-B (Stemcell Technologies 07156), 0.5mL 55mM β-mercaptoethanol (Life technologies 21985-023), 0.5mL primocin (Invivogen ant-pm-2), BDNF (10ng/mL, peprotech 450-02), NT-3 (10ng/mL, peprotech 450-03), laminin (200ng/mL, Sigma L2020), and doxycycline (2ug/mL, Sigma D3072) in DMEM / F12 (Gibco 11320-033) for 24 hours (DIV0). Cells were then exposed to puromycin (5ug/mL, Sigma P9620) for 24 hours in the same N2 media (DIV1). iNeurons were re-plated 24 hours later (DIV2) to experiment appropriate plates coated with Matrigel GFR (Corning 354230). Cells were washed 2x in PBS and lifted with StemPro accutase (Thermo Fisher Scientific A11105-01) for 5min at 37°C. Cells were spun down at 1000RPM for 5min at RT. Cells were resuspended and plated in iN media (Neurobasal-A media (Invitrogen A24775-01) with 5mM glucose (Sigma G5146), 10mM sodium pyruvate (Sigma P5280), glutamax (Life Technologies 35050-061), penicillin/streptomycin (Thermo Fisher Scientific 15140-148), BDNF and NT-3 (10ng/mL), and doxycycline (2ug/mL) through 9 days. 2µM Ara-C (Sigma C6645) was added on DIV3, and there was a full media exchange 24 hours later (DIV4). Doxycycline post DIV10 was discontinued for the rest of the 21-day differentiation.

### iAstrocyte differentiation of iPSCs

iPSCs transfected with the NGN2 virus were put through the first two days of the iNeuron differentiation. On day 3, cells were exposed to Astrocyte differentiation media: N2 media (without BDNF, NT-3, laminin, and doxycycline), 10% FBS (Hyclone SH30071.03HI), B-27 with vitamin-A (Thermo Fisher Scientific 17504044), FGF-2 (20ng/mL, R&D Systems 233-FB-025) and EGF (20ng/mL, R&D Systems 236-EG-200). After 30 days, EGF was removed and FGF2 reduced to 5ng/mL. After 70 total days, cells were switched into Astrocyte Media (Sciencell 1801). Experiments were performed after 90 total days of differentiation. After DIV90, iAstrocyte stocks were frozen down.

### iMicroglia differentiation of iPSCs

iPSCs were differentiated into common myeloid progenitors (CMPs) according to the published protocol (Paluru et al., 2014) by the Human Pluripotent Stem Cell Core (CHOP). CMPs were plated at 333K cells / well in a 24-well Cellbind plate (Corning 3337). CMPs were differentiated in iMg media (RPMI 1640 media (GE Healthcare Life Sciences SH30027.01) with 10% FBS (Hyclone SH30071.03HI), Recombinant Human IL-34 (100ng/mL, R&D systems 5265-IL-010), CSF-1 Recombinant Human Protein (25ng/mL, Thermo Fisher Scientific PHC9504), and Recombinant Human TGF-β1 (50ng/mL, Peprotech 100-21). Half media changes were performed every two days for 11 days.

### Tri-culture combination

iNeurons were differentiated as described above and re-plated on DIV2 to Matrigel (Corning 354230) coated Nunc Lab-Tek II 8-well chamber slides (Thermo Fisher Scientific 62407-296) at 70k cells/well in iN media. On DIV5 of iNeuron differentiation, the iAstrocytes were added at 50k cells/well. On DIV7 of iNeuron differentiation, iMicroglia were added at 100k cell/well. Cultures were taken out to DIV21 of iNeuron differentiation. Of note, this preparation creates a “carpet-culture” of the three cell types that is roughly 50um thick

### Human astrocytes

Three separate donors for human astrocytes were obtained from Sciencell. Cells were plated on PLL-coated plates (Sigma Aldrich P6282) and grown for two weeks in Astrocyte media (Sciencell 1801) and then RNA was extracted.

### iAstrocyte cytokine exposure

iAstrocytes in mono-culture were exposed to IL-1b (R&D Systems 201-LB-005) or IL-8 (R&D Systems 208-IL-010) at 10ng/mL or PBS vehicle control for 8 hours. Supernatants were collected and sent for cytokine analysis.

### HIV infection

iMicroglia were differentiated to D11. On D11, they were exposed to 50ng/mL of JAGO strain HIV (UPenn Center of Aids Research; CFAR) for 24 hours. Media was fully exchange after 24 hours and collected. Full media exchanges occur every 3 days for 15 days. If antiretroviral treatment was used, it was started on day 6 of infection and added with each media exchange. Efavirenz (U.S. Pharmacopeia 1234103) was used at 20nM and Darunavir (Prezista TMC114) at 4.5µM.

### Monocyte-derived macrophage differentiation

We receive donated buffy coat from New York Blood Center from three donors (D471, D446, D470). Buffy coat was diluted 1:1 with PBS (without Ca^2+^ and Mg^2+^) (Invitrogen 14190144). 15 mLs of Ficoll (Sigma Aldrich 26878-85-8) was added to 50mL conical tubes. 25mLs of Buffy Coat/PBS was slowly layered onto the Ficoll. Samples were spun at 1200 rpm for 45 minutes with no brake. The peripheral blood mononuclear cell (PBMC) layer was removed and placed into a new 50mL conical tube, and the volume was brought up to 50mL with PBS. The PBMCs are spun at 450Xg for 10 minutes. The supernatant was discarded, and the pellet resuspended in 10mLs of Red Blood Cell Lysis buffer (Sigma Aldrich 11814389001). Cells were shaking at RT for 10 minutes. Volume was brought up to 50mLs in PBS and spun at 450Xg for 10minutes. The supernatant was discarded, and the pellet was resuspended in DMEM with 10% FBS + Gentamicin (Thermo Fisher Scientific 15750060). The PBMCs were plated on 6-well tissue culture plate for 5 days. On day 5, a full media exchange was performed and added 10ng/mL Human GM-CSF (Gold Biotechnology 1120-03-20). A half media exchange was performed on day 7. At day 10, RNA was collected from macrophages.

### RNA extraction

Cells were washed twice with RT PBS, then lifted with StemPro accutase (Thermo Fisher Scientific A11105-01) and spun down at 1,500 rpm for 5 min. Fresh cells were processed through Qiashredder (Qiagen 79654) and RNeasy mini kit (Qiagen 74104) and frozen at −80°C.

### Immunofluorescence

Cells were washed twice with PBS, then fixed in 4% PFA (VWR TCP0018) for 15 min at RT. Cells were washed 3x in PBS for 5 min at RT before being stored in PBS at 4°C. Cells are blocked in 5% BSA (Sigma A9418) and 0.1% Triton X-100 (sigma X100) for 1 hr at RT. Sections use 0.3% Triton-X 100 in blocking buffer. Primary and secondary antibodies were diluted in blocking buffer. Cells were incubated in primary antibodies overnight at 4°C. Cells were washed 3x 5min each at RT in PBS-T (0.1% Tween20 (Sigma P9416). Secondary was performed at RT in the dark for 1 hour. Cells were washed 3x 5min each at RT in PBS-T, then mounted with Prolong gold antifade (Life Technologies P36930).

Antibodies used for immunofluorescence: Chicken anti-MAP2 (Abcam ab5392, RRID:AB_2138153, 1:500); Mouse anti-PSD-95 clone K28/43 (NeuroMab 75-028, RRID:AB_2292909, 1:500); Mouse anti-Synaptophysin clone SY38 (Millipore MAB5258-20UG, RRID:AB_95185, 1:250); Mouse anti-Nestin clone 10C2 (Millipore MAB5326, RRID:AB_11211837, 1:200); Rabbit anti-Thrombospondin-1 (Abcam ab85762, RRID:AB_10674322, 1:250); Mouse anti-Glutamine Synthetase (Millipore MAB302, RRID:AB_2110656, 1:500); Rabbit anti-SOX9 (Abcam ab185230, RRID:AB_2715497, 1:250); Rabbit anti-CX3CR1 (Abcam ab8021, RRID:AB_306203, 1:500); Rabbit anti-TMEM119 (Abcam ab185333, RRID:AB_2687894, 1:200); Rabbit anti-IBA1 (Wako 019-19741, RRID:AB_839504, 1:500); Rabbit anti-P2RY12 (Alomone Labs APR-020, RRID:AB_11121048, 1:100); Rat anti-LAMP1 (Abcam ab25245, RRID:AB_449893, 1:500); Mouse anti-HIV1 p24 [39/5.4A] (Abcam ab9071, RRID:AB_306981, 1:500); Rabbit anti-Human Nanog (Cell Signaling Technology 3580, RRID:AB_2150399, 1:800); Mouse anti-NCAM Clone 2-2b (Millipore MAB5324, RRID:AB_95211, 1:250); Rabbit anti-OCT-4 (Cell Signaling Technology 2750, RRID:AB_823583, 1:200); Rabbit anti-SOX2 (Millipore AB5603, RRID:AB_2286686, 1:100); Rabbit anti-CCR5 (Thermo Fisher Scientific pa5-29011, RRID:AB_2546487, 1:500); Mouse anti-GLAST (Miltenyi Biotec 130-095-822, RRID:AB_10829302, 1:50); Mouse anti-GFAP (Sigma-Aldrich SAB1405864, RRID:AB_10739114, 1:10,000); Mouse anti-Beta-Actin (Cell Signaling Technology 3700, RRID:AB_2242334, 1:10,000); Rabbit anti-EAAC1 (Santa Cruz Biotechnology sc-25658, RRID:AB_2190727, 1:50); Rabbit GLT-1 (Jeff Rothstein Lab 1:5,000); Goat anti-mouse IgG (H+L) Alexa Fluor 488 (Thermo Fisher Scientific A-11029, RRID:AB_138404, 1:500); Goat anti-rabbit IgG (H+L) Alexa Fluor 488 (Thermo Fisher Scientific A-11034, RRID:AB_2576217, 1:500); Goat anti-mouse IgG (H+L) Alexa Fluor 568 (Thermo Fisher Scientific A-11004, RRID:AB_2534072, 1:500); Goat anti-rabbit IgG (H+L) Alexa Fluor 568 (Thermo Fisher Scientific A-11036, RRID:AB_10563566, 1:500); Goat anti-rat IgG (H+L) Alexa Fluor 568 (Molecular Probes A-11077, RRID:AB_141874, 1:500); Goat anti-rat IgG (H+L) Alexa Fluor 680 (Molecular Probes A-21096, RRID:AB_141554, 1:500); Goat anti-Chicken IgY (H+L) DyLight 680 (Thermo Fisher Scientific SA5-10074, RRID:AB_2556654, 1:500)

### iAstrocyte immunofluorescence counting

Images were obtained with a Nikon eclipse N1 scope equipped with LED-based epifluorescence. The optical fractionator workflow mode of Stereo Investigator 64 bit was then used to generate random areas of the wells to image. 5 images were obtained per well. Images were then transferred to image J where the channels were manually merged. After merging images, the “cell count” plug in of imageJ is used to quantify the total number of DAPI (+) cells, and the number of co-labeled DAPI+ astrocyte marker(+) cells.

### Local Synapse Density

Images were analyzed on ImageJ. The iMicroglia were outlined and had a 50µm radius circle drawn from the center of the cell with ROI manager. The total synapse density was measured using the analyze particles plugin. The plugin was used for the ROI of the cell and the surrounding circle. The area of the cell and the area of the particles were subtracted from the total area of the circle and the total particles. The remaining total area of the particles was divided by the remaining total area of the circle to calculate local synapse density.

### Bulk RNA-seq

RNA was extracted using previously described with RNeasy mini kit (Qiagen 74104) and frozen at −80°C. Samples were sent to the Center for Applied Genomics (CAG) for sequencing. In short, the Illumina TruSeq Stranded Total RNA library kit (Illumina RS-122-2201) for RNA-seq was utilized for preparation of the libraries for sequencing. Libraries were produced using liquid handler automation with the PerkinElmer Sciclone instrument. This procedure started with a ribosomal RNA (rRNA) depletion step. The depletion step uses target-specific oligos with specialized rRNA removal beads to remove both cytoplasmic and mitochondrial rRNA from the total RNA. Following this purification, the RNA was fragmented using a brief, high-temperature incubation. The fragmented RNA was then reverse transcribed into first-strand cDNA using reverse transcriptase and random primers. Second strand cDNA was generated using DNA Polymerase which was then used in a standard TruSeq Illumina-adapter based library preparation. Library preparation consisted of four main steps: unique adapter-indexes were ligated to the RNA fragments, AmpureXP bead purification occurred to remove small library fragments, the libraries were enriched and amplified using PCR, and the libraries underwent a final purification using AmpureXP beads. Upon completion, library quality was assessed using an automated electrophoresis instrument, the PerkinElmer Labchip GX Touch, and qPCR using the Kapa Library Quantification Kit and Viia7 real-time PCR instrument. Libraries were diluted to the appropriate sequencer loading concentration and pooled accordingly to allow for the desired sequencing depth. RNA libraries were sequenced in one lane of the Illumina HiSeq2500 sequencer using the High Output v4 chemistry and paired-end sequencing (2×100bp).

### Single Cell RNA-seq cell preparation

Cells were incubated in 0.25% Trypsin+EDTA at 37°C for 8 minutes, put through a cell strainer, and spun down in ice-cold PBS at 4°C for 5 min at 1500 RPM. Cells were resuspended in ice-cold DPBS without Mg^2+^ and Ca^2+^+0.04% BSA. Up to 20,000 cells were sent for sequencing per sample. Samples were sent to CAG for sequencing.

### Bulk RNA-seq analysis

RNA-seq reads were demultiplexed using the DRAGEN genome pipeline (Goyal et al., 2017). FASTQ files were aligned to hg19 reference using the STAR (v.2.6.1) (Dobin et al., 2013) aligner with default settings. Generated BAM files were read into the R statistical computing environment. Gene counts were obtained using the GenomicAlignments package. Differential expression analysis was performed using the R/Bioconductor package DESeq2 which uses a negative binomial model (Love, Huber, & Anders, 2014). Analysis was performed using standard parameters with the independent filtering function enabled to filter genes with low mean normalized counts. The Benjamini-Hochberg adjustment was used to estimate the false discovery rate (Padj) and correct for multiple testing. Genes were then analyzed using the Ingenuity IPA software (QIAGEN Inc.). Additional published RNA-seq data was utilized for comparative analysis from project accession SRP092075 (Abud et al., 2017). Datasets were obtained and converted to fastq format using the Sequence Read Archive (SRA) tool provided by NCBI. Fastq-formatted data was analyzed similarly to the bulk RNA-seq samples using the DRAGEN pipeline and integrated into the experimental R data object.

### Single Cell RNA-seq analysis

Next-generation sequencing libraries were prepared using the 10x Genomics Chromium Single Cell 3’ Reagent kit v2 per manufacturer’s instructions. Libraries were uniquely indexed using the Chromium i7 Sample Index Kit, pooled, and sequenced on an Illumina Hiseq sequencer in a paired-end, single indexing run. Sequencing for each library targeted 20,000 mean reads per cell. We had a mean of 39,227 reads per cell post-normalization with 2,165 median genes per cell. Data was then processed using the Cellranger pipeline (10x genomics, v.3.0.2) for demultiplexing and alignment of sequencing reads to the GRCh38 transcriptome and creation of feature-barcode matrices. Individual single cell RNAseq libraries were aggregated using the cellranger aggr pipeline. Libraries were normalized for sequencing depth across all libraries during aggregation.

Secondary analysis on the aggregated feature barcode matrix was performed using the Seurat package (v.3.0) within the R computing environment. Briefly, cells expressing less than 200 or more than 5000 genes were excluded from further analysis. Additionally, cells expressing >20% mitochondrial genes were excluded from the dataset. Log normalization and scaling of features in the dataset was performed prior to principal component dimensionality reduction, clustering, and visualization using tSNE. Cell types were identified using expression of canonical cell markers in microglia (AIF1, SPI1, CD4), neurons (MAP2, SYN1), and astrocytes (THBS1, SOX9). Differentially expressed genes and identification of cluster or cell type specific markers were identified using a wilcoxon rank sum test between two groups. P-value adjustment was performed using bonferroni correction based on total number of genes in the aggregated dataset. Genes were then analyzed using the Ingenuity IPA software (QIAGEN Inc.).

### qRT-PCR

RNA was extracted by RNeasy mini kit (Qiagen 74104). cDNA was generated using SuperScript VILO Master mix (Thermo Fisher Scientific 11755050). RNA expression was measured using Taqman probes (Thermo Fisher Scientific) for *CX3CR1* (Hs01922583_s1), *P2RY12* (Hs01881698_s1), *TMEM119* (Hs01938722_u1), *THBS1* (Hs00962908_m1), and *GAPDH* (Hs02786624_g1). 30ng of cDNA was used per well, with three technical replicates per probe per sample. qRT-PCR is run on an Applied Biosystems 7900HT Fast Real-Time PCR System. All expression levels were normalized to *GAPDH* expression.

### Reverse transcriptase activity

10 µl per well of supernatant was placed into a 96-well microtiter plate to be analyzed for RT activity. 50 µl of RT cocktail was added per well into the 96-well microtiter plate. RT cocktail consists of: 50mM Tris (Amresco J837) pH 7.8, 75mM KCl (Ambion 9610), 2mM Dithiothreitol (DTT) (Sigma D0632), 5mM Magnesium Chloride (Ambion 9530G), 5ug/mL Polyadenylic acid (GE Healthcare 27-4110-01), 2.5ug/mL pd (t)12-18 (Oligo dT) (USB Corporation #19817), 0.05% NP-40 (Calbiochem, 492016), 10 µCi/ml Thymidine 5’-triphosphate, ALPHA-[32P] / [32P] TTP (Perkin Elmer BLU005A250MC). Samples were incubated at 37° C overnight. 30 µl of RT reaction mixture was placed onto pre-marked DE81 paper (Whatman 3658915) and air dried for 15 min at RT. Paper was washed 4x, 5 min each with 2x SSC (Roche-Apply science 11 666 681 001) by submerging in a tray on a rotating platform. Paper was then washed 1x, 1 min in 100% ethanol and air dried in an oven at 80-100° C (25-30 min). Each paper sample was placed into a scintillation vial. 5 ml Betaflour (National Diagnostics LS-151) was added to each scintillation vial. 32P was counted on a scintillation counter, yielding CPM.

### PCR for CCR5Δ32 mutation

DNA was isolated with DNeasy blood and tissue kit (Qiagen 69504). DNA Oligos were generated by IDT. Forward primer: 5’ – CAAAAAGAAGGTCTTCATTACACC – 3’. Reverse primer: 5’ – CCTGTGCCTCTTCTTCTCATTTCG – 3’. Primers were reconstituted to 100uM. Mastermix consists of 10uL KAPA PCR buffer, 7uL H20, 1uL 10uM forward primer, 1uL 10uM reverse primer. 1uL DNA added separately. 1.5% agarose gel was run for 1hour at 120v. The gel was imaged on a Biorad Universal Hood II Gel Doc System.

### Western Blot

iAst plated in 12-well dishes were rinsed twice with PBS containing 0.1 mM Ca^2+^ and 1.0 mM Mg^2+^ (PBS Ca^2+^/Mg^2+^) then lysed in 200ul of radioimmunoprecipitation (RIPA) buffer (100 mM Tris-HCl, pH 7.4, 150 mM NaCl, 1 mM EDTA, 0.1% SDS, 1% Triton X-100, 1% sodium deoxycholate containing protease inhibitors, including 1 mg/ml leupeptin, 250 mM phenylmethylsulfonyl fluoride, 1 mg/ml aprotinin, and 1 mM iodoacetamide) for 1 hour while rotating on a shaker at 4°C. Cortical and hippocampal tissue was harvested from adult C57BL/6 mice and solubilized in 5 volumes of homogenization buffer (50 mM Tris-HCl, pH 7.4, 150 mM NaCl, 5 mM EDTA, 1% NP-40, 1% SDS, containing protease and phosphatase inhibitors, including 1 mg/ml leupeptin, 250 mM phenylmethylsulfonyl fluoride, 1 mg/ml aprotinin, 1 mM iodoacetamide, 10mM NaF, 30mM Na pyrophosphate, 1mM Na^3^VO^4^). All lysates were centrifuged at 17,000g for 20 min to remove cellular debris and nuclei. The supernatants were analyzed for total protein using the Pierce protein assay kit according to the manufacturer’s instructions. Lysates are diluted 1:1 with 2X Laemmli buffer and either boiled at 95°C for 5 min (GFAP, Actin) or kept at 25°C for 45 minutes (GLAST, GLT-1, EAAC1). iAst lysates (20ug) or cortical/hippocampal lysates (5ug) were then resolved by SDS PAGE using 10% BioRad minigels and transferred to Immobilon-FL membranes (Millipore Cat# IPFL00010). Membranes were incubated with blocking buffer (1% non-fat dry milk in TBS-T) for 1 hour at 25°C prior to probing with primary antibodies diluted in blocking buffer overnight at 4°C: rabbit anti-GLT-1 (C-terminal directed 1:5,000; Rothstein et al 1994), mouse anti-GLAST (Miltenyi Biotec Cat# 130-095-822, 1:50), rabbit anti-EAAC1 (Santa Cruz Cat# SC-25658, 1:50), mouse anti-GFAP (Cell Signaling Cat#3670S, 1:1,000), and mouse anti-Actin (1:10,000 dilution, Cell Signaling, Cat# 3700S). Membranes were washed with blocking buffer 3 times for 10 min at 25°C. After the washes, membranes were probed with anti-mouse or anti-rabbit fluorescently conjugated secondary antibodies (LI-COR Biosystems, 1:10,000) for 45 minutes at 25°C. The membranes were washed 3 times for 10 minutes each then visualized using a LI-COR Odyssey.

### Synaptophagocytosis analysis

Confocal images of microglia were analyzed in IMARIS software. IBA1 labeling for the microglia was surfaced, as well as synaptophysin staining. Total volume was taken for the microglia and the synaptophysin staining inside the cells. Total synaptophysin volume per cell was then divided by the cell volume defined by IBA1 labeling.

### Cytokine analysis

Supernatants were tested for 6 analytes (TNFa, IL-6, IL-8, IL-10, IL-1b, IL-1a) on a custom Human magnetic Luminex plate (R&D systems LXSAHM; run by Penn Mental Health AIDS Research Center (PMHARC)). The plate was run by the Penn Mental Health Aids Research Center on a MAGPIX powered by Luminex XMAP technology.

## Acknowledgements

This work was supported by the National institute of Neurological Disorders and Stroke (R21 NS107594 02) and the Penn Center for Aids Research (CFAR) and Penn Mental Health AIDS Research Center (PMHARC) (5-P30-MH-097488-05). We would like to thank the Human Pluripotent Stem Cell Core (CHOP) for providing CMPs, Center for Applied Genomics for performing and analyzing RNAseq, CFAR for providing HIV strains, and PMHARC for analyzing cytokines. We would like to thank Kieona Cook for image analysis and Elizabeth Krizman for performing western blots, who was partially supported by the Intellectual and developmental Disabilities Research Center at CHOP/Penn U54 HD086984, and Herbert M. Lachman, MD for providing iPSC lines.

