## Supplemental figures for "Neuroinflammation and EIF2 signaling persist in an HiPSC tri-culture model of HIV infection despite antiretroviral treatment"

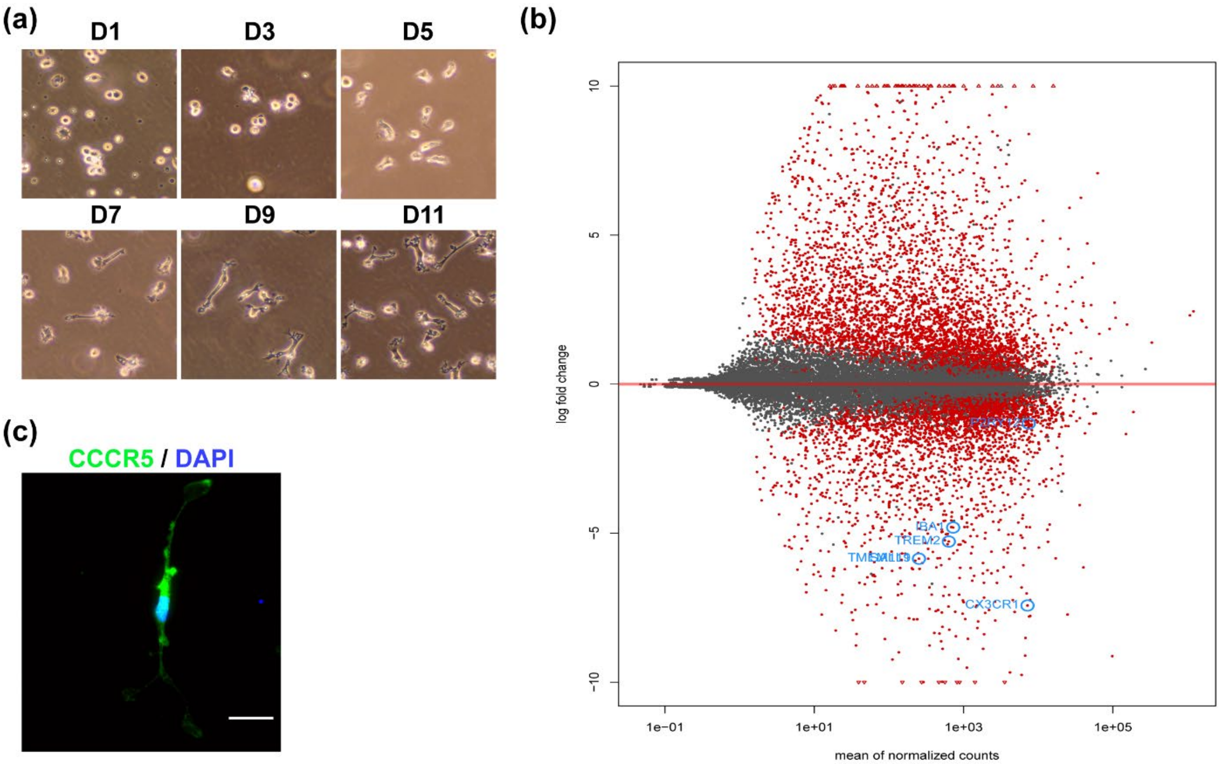


**Figure 1-figure supplement 1: Morphology and comparative RNA expression of iMG to MDMs**

(a) Brightfield images of iMg differentiation, depicting ramification by D11. (b) MA plot of bulk RNAseq data comparing iMg (baseline) to MDMs. n=3 Benjamini-Hochberg FDR-0.05. (c) Immunostaining showing CCR5 expression (green) on D11 iMg. scale bar represents 25um.

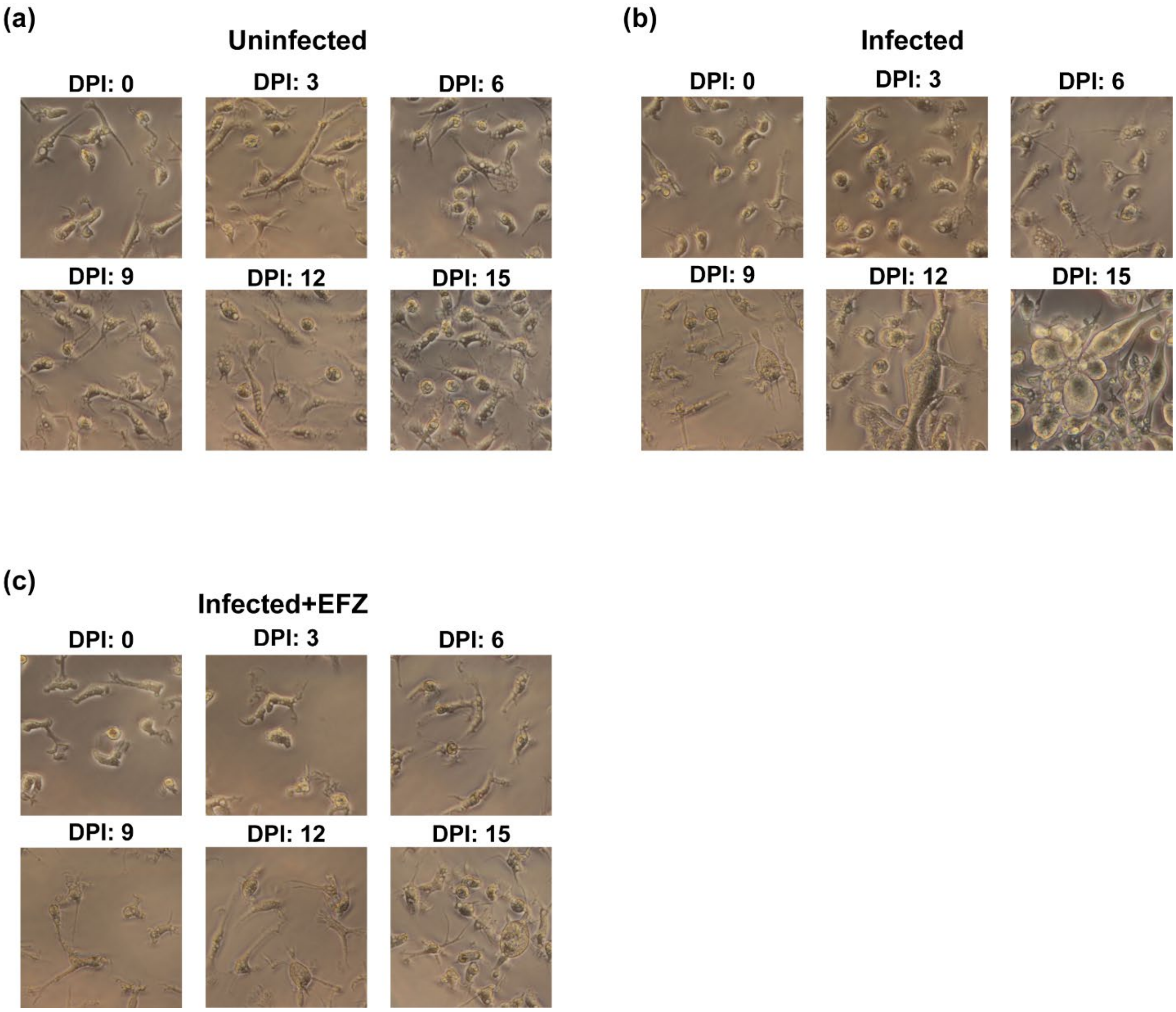


**Figure 2-figure supplement 1: Morphology changes during HIV infection ± EFZ**

(a) Brightfield images over infection period in an uninfected culture shows no syncytia. (b) Brightfield images over infection period in an infected culture shows syncytia beginning at D6-9 and most cells by D15. (c) Brightfield images over infection period in an Inf+EFZ culture shows a reduced rate of syncytia compared to infected.

(a) (b)

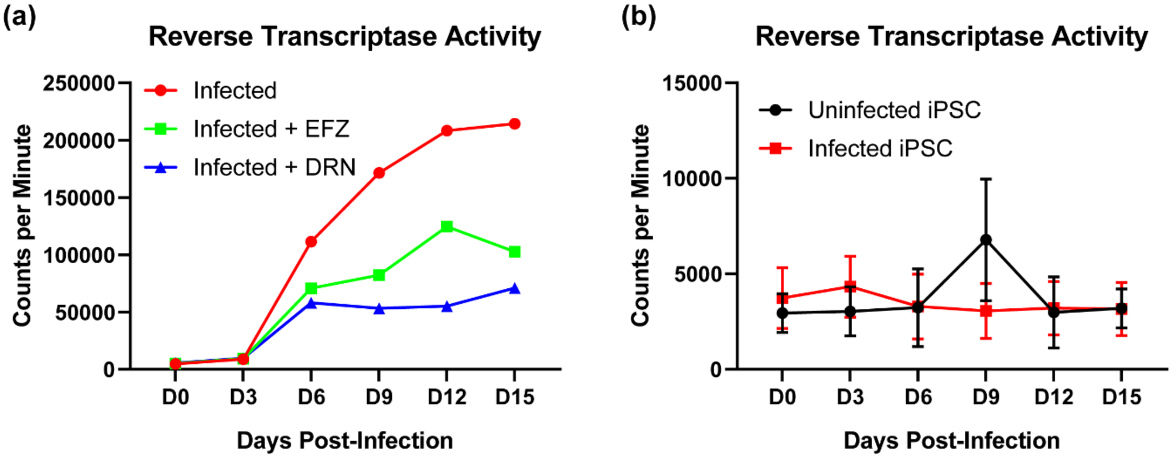


(c)

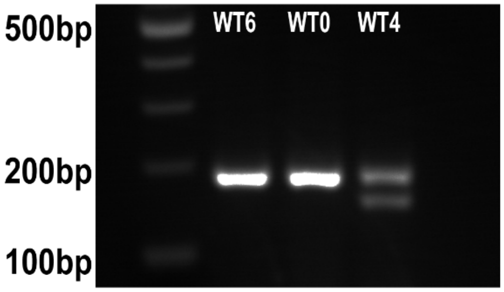


(d)

| Cell Line | Number of P24 Reads |
|---|---|
| WT6+HIV | 3759 |
| WT10+HIV | 3535 |
| WT15+HIV | 1876 |
| WT4+HIV (CCR5Δ32 Het) | 150 |
| WT6 | 0 |
| WT10 | 0 |
| WT15 | 0 |
| WT4 | 0 |

**Figure 2-figure supplement 2: Extended characterization of infected ± ART iMg in mono-culture and CCR5Δ32 mutation**

(a) Reverse Transcriptase activity of WT6 exposed to EFZ (20nM) or DRN (4.5µM) starting D6 confirms iMg respond well to antiretroviral treatment. n=1 infection. (b) Reverse transcriptase activity of iPSCs exposed to 50ng/mL JAGO shows iPSCS were unable to be infected at the virus concentration used for the iMg. n=3 infections, one-way ANOVA Dunnett's post hoc analysis, error bars represent SEM. (c) PCR of CCR5Δ32 shows WT4 is heterozygous for the mutation. (d) Infected WT4 has dramatically less raw reads of P24 from Bulk RNAseq compared to infected WT6 and WT10 iMg.

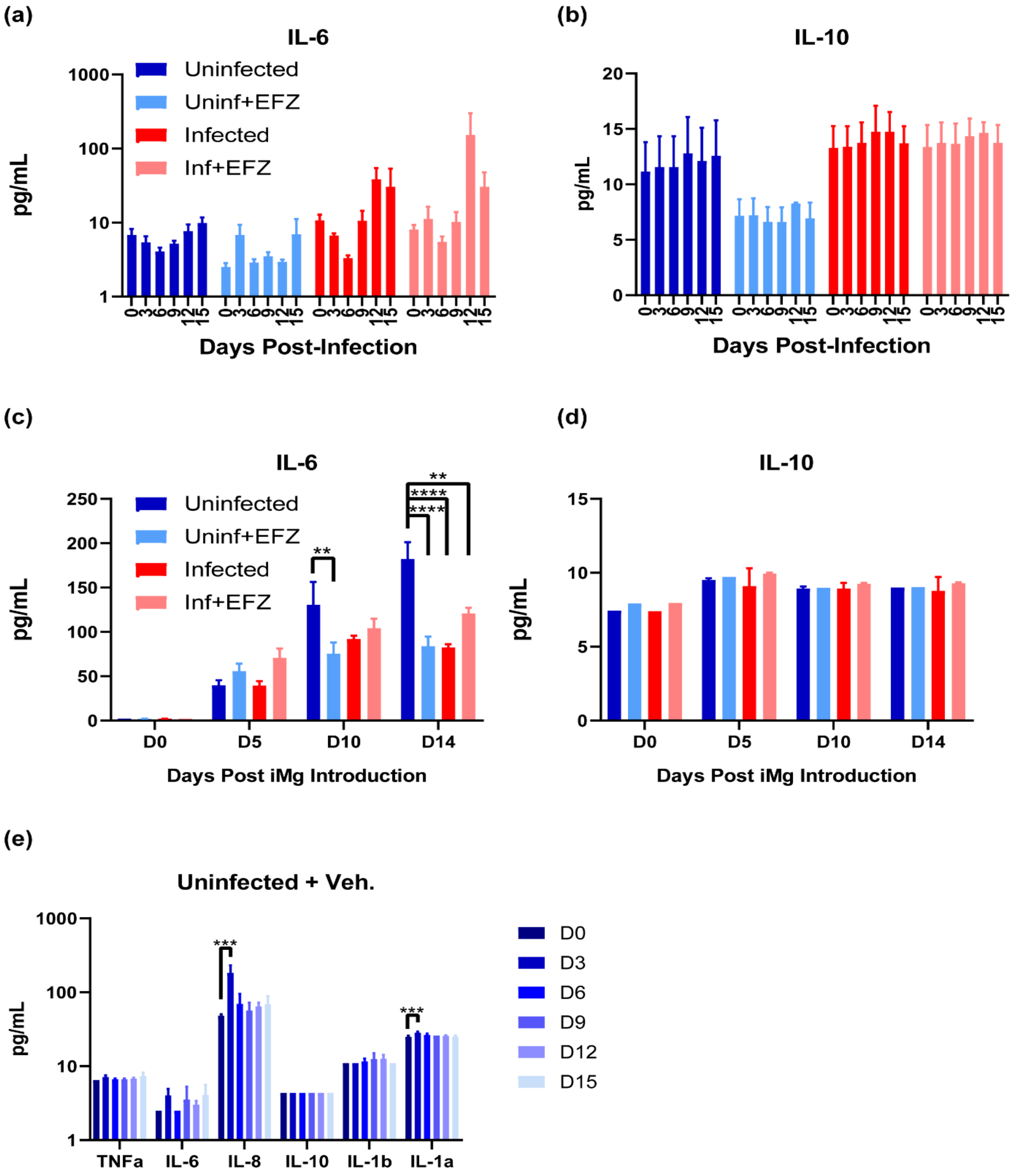


**Figure 2-figure supplement 3: Additional cytokine expression during infection and with vehicle control**

(a&b) Il-6 (a) and IL-10 (b) production in mono-culture did not change for any of the four conditions. n=4 Uninf and Uninf+EFZ, n=3 Inf and Inf+EFZ, one-way ANOVA, Dunnett's post hoc analysis, error bars represent SEM. (c&d) Tri-culture Il-6 (c) production increased over time for Uninf, and no change in IL-10 (d) for any of the four conditions. n=3 differentiations, two-way ANOVA, Tukey's post hoc analysis, **p<0.01; ****p<0.0001, error bars represent SEM. (e) Uninf+ Veh. had minimal but significant increases in IL-8 and IL-1a production at D3, but no increase at D12 where infected cultures had significant increases. n=3 differentiations, one-way ANOVA, Dunnett's post hoc analysis, ***p<0.001, error bars represent SEM.

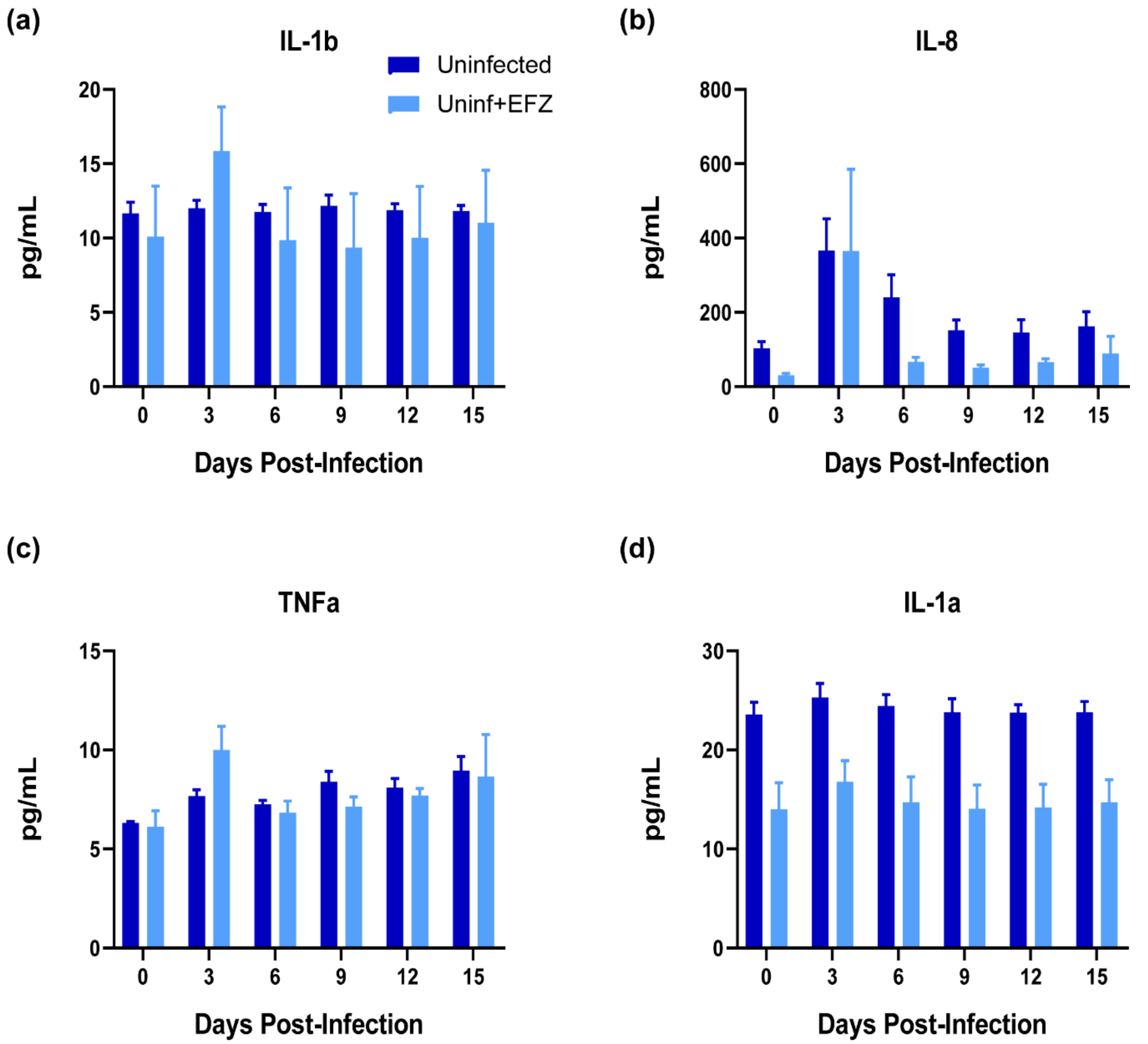


**Figure 2-figure supplement 4: No change in IL-1b, IL-8, TNFa, or IL-1a in Uninf and Uninf+EFZ mono-culture iMg**

(a-d) cytokine production in Uninf and Uninf+EFZ mono-cultures. Il-1b (a), IL-8 (b), TNFa (c), and IL-1a (d) production in mono-culture did not change for Uninf or Uninf+EFZ. n=4 differentiations, one-way ANOVA, Dunnett's post hoc analysis, error bars represent SEM.

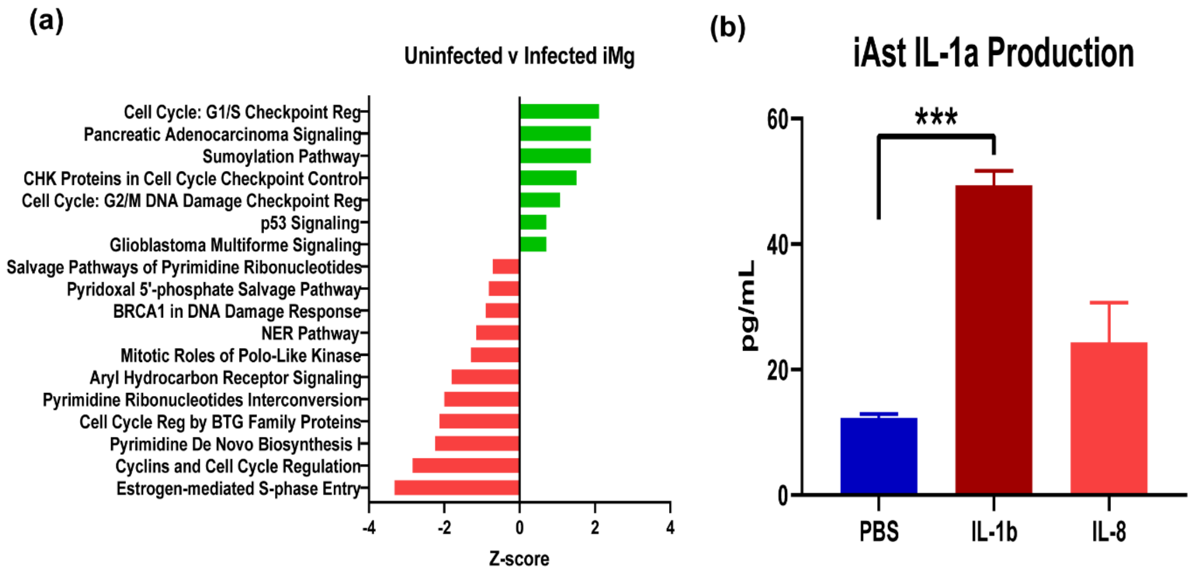


(c)

| Gene | log2fold change | FDR |
|---|---|---|
| *CCL8* | 4.425125 | 0.026731 |
| *C5AR1* | 1.249691 | 0.026599 |
| *C2* | 1.305198 | 4.56E-05 |
| *CD80* | 3.148555 | 3.43E-05 |
| *NFKBIZ* | 1.866165 | 0.003559 |
| *NFKBID* | 1.53444 | 0.000622 |
| *FOS* | 2.185966 | 0.003814 |
| *IL1B* | 2.164145 | 0.001461 |
| *PTGS2* | 3.682552 | 0.002386 |
| *ATF3* | 2.755115 | 9.45E-07 |
| *CXCR2* | 1.056328 | 0.02422 |
| *TNFRSF11A* | 2.065118 | 0.009194 |
| *TNFRSF10B* | 1.299788 | 0.00607 |

**Figure 2-figure supplement 5: Bulk RNAseq analysis of uninfected versus infected iMg in mono-culture and IL-1a production in iAst**

(a) Ingenuity Pathway Analysis of Uninfected v Infected iMg bulk RNAseq. n=3 cell lines, Fisher's exact <0.05. Benjamini-Hochberg FDR 0.05. (b) iAst produced IL-1a in response to 8hr exposure to IL-1b (10ng/mL), but not IL-8 (10ng/mL). n=3 cell lines, one-way ANOVA, Dunnett's post hoc analysis, ****p<0.0001, error bars represent SEM. (c) Select inflammation related genes upregulated in iMg+HIV bulk RNA-seq. Log2fold change at least 1 and FDR <0.05.

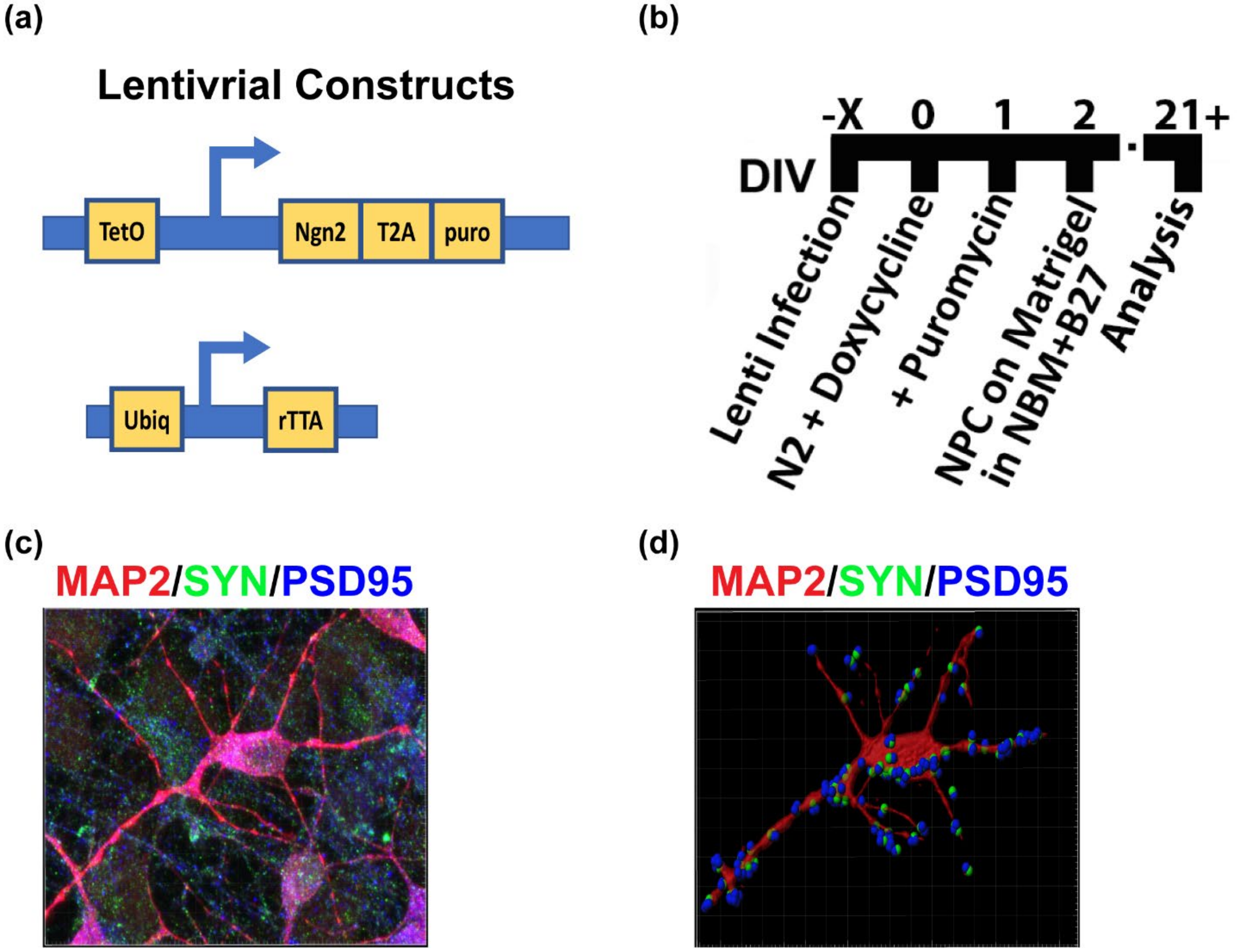


**Figure 3-figure supplement 1: Differentiation timeline and validation of iNrn**

(a) schematic of the two lentiviral vectors for the iNrn differentiation. (b) Timeline for iNrn differentiation. (c) Immunostaining and surface reconstruction of MAP2+ (red) iNrn with apposing pre- and post-synaptic puncta, synaptophysin (green) and PSD-95 (blue).

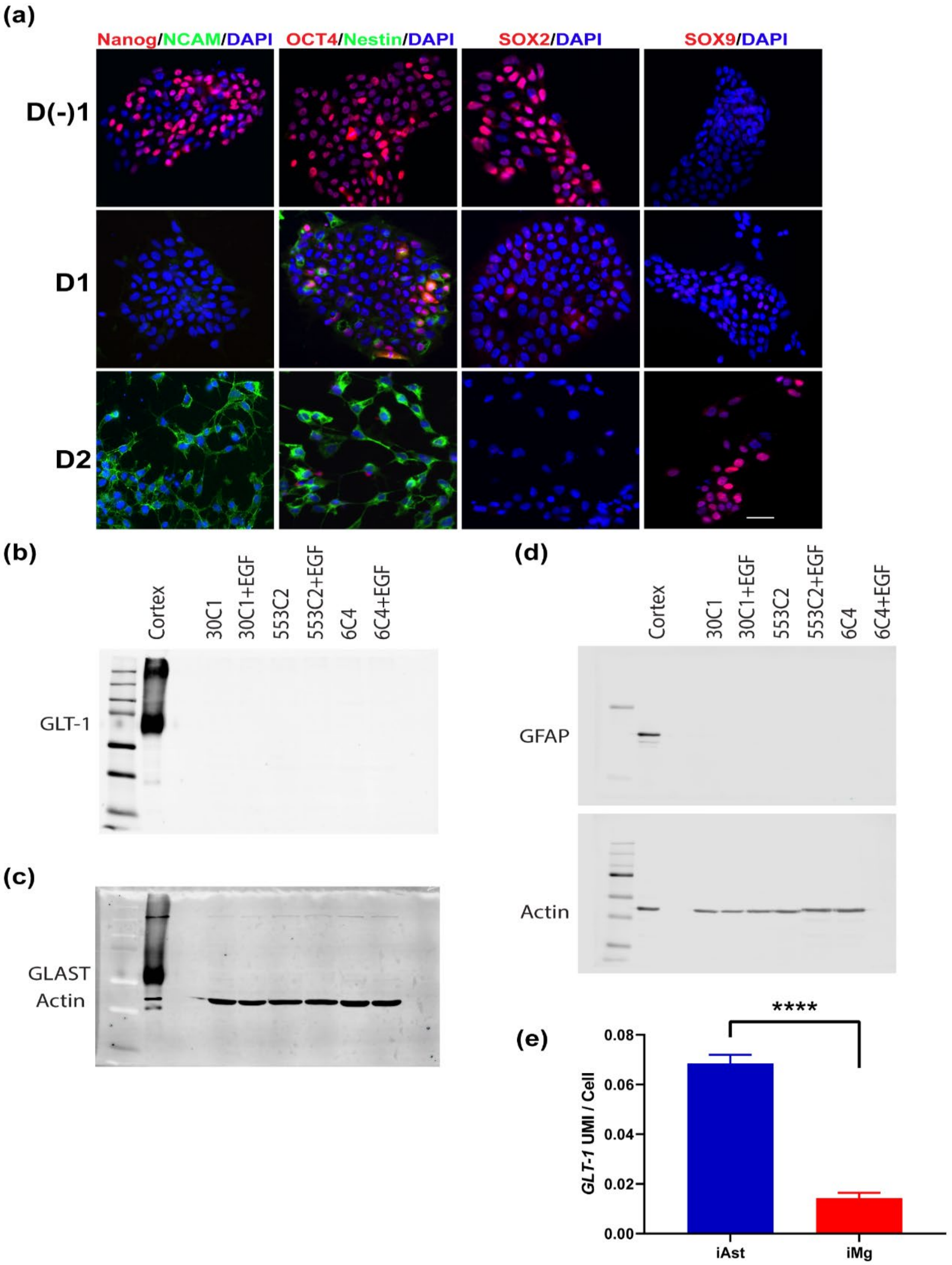


**Figure 3-figure supplement 2: NPC markers during early iNrn differentiation and *GLT-1* expression in tri-culture**

(a) By day 2 of the NGN2 differentiation (post puromycin selection), cells express have lost pluripotency markers (Nanog, OCT4, and Sox2) and begin to express neural progenitor markers Nestin and NCAM, as well as, the astrocyte marker Sox9. Scale bar represents 25µm. (b-d) Western blot showing iAst do not express glutamate transporter GLT-1 (b), GLAST (c), or GFAP (d) in mono-culture. (e) scRNAseq analysis shows that, in tri-culture, iAst express *GLT-1*. iAst n=4,763; iMg n=4,485. two-tailed t test, ****$p<0.0001$, error bars represent SEM.

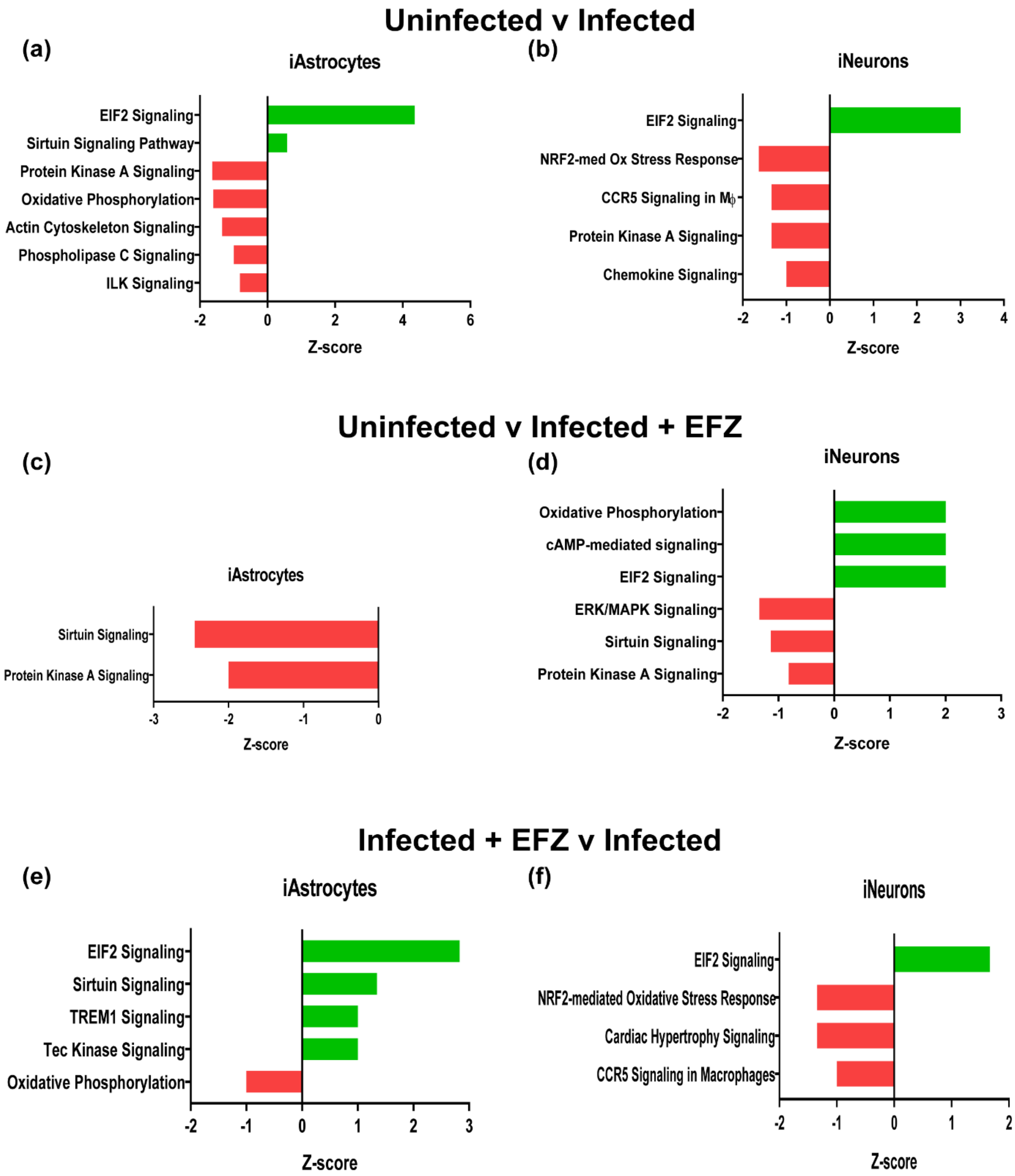


**Figure 6-figure supplement 1: iAst and iNrn Ingenuity pathway analysis for Uninf v Inf, Uninf v Inf+EFZ, and Inf+EFZ v Inf**

(a&b) Top affected pathways from Ingenuity pathway analysis of iAst (a) and iNrn (b) in the uninfected versus infected conditions. Uninfected is baseline. Benjamini-Hochberg FDR 0.05, Fisher's exact <0.05, z-score cutoff ± 0.5. (c&d) Top affected pathways from Ingenuity pathway analysis of iAst (c) and iNrn (d) in the uninfected versus infected + EFZ conditions. Uninfected is baseline. Benjamini-Hochberg FDR 0.05, Fisher's exact <0.05, z-score cutoff ± 0.5. (e&f) Top affected pathways from Ingenuity pathway analysis of iAst (e) and iNrn (f) in the Infected + EFZ versus infected conditions. Infected + EFZ is baseline. Benjamini-Hochberg FDR 0.05, Fisher's exact <0.05, z-score cutoff ± 0.5.

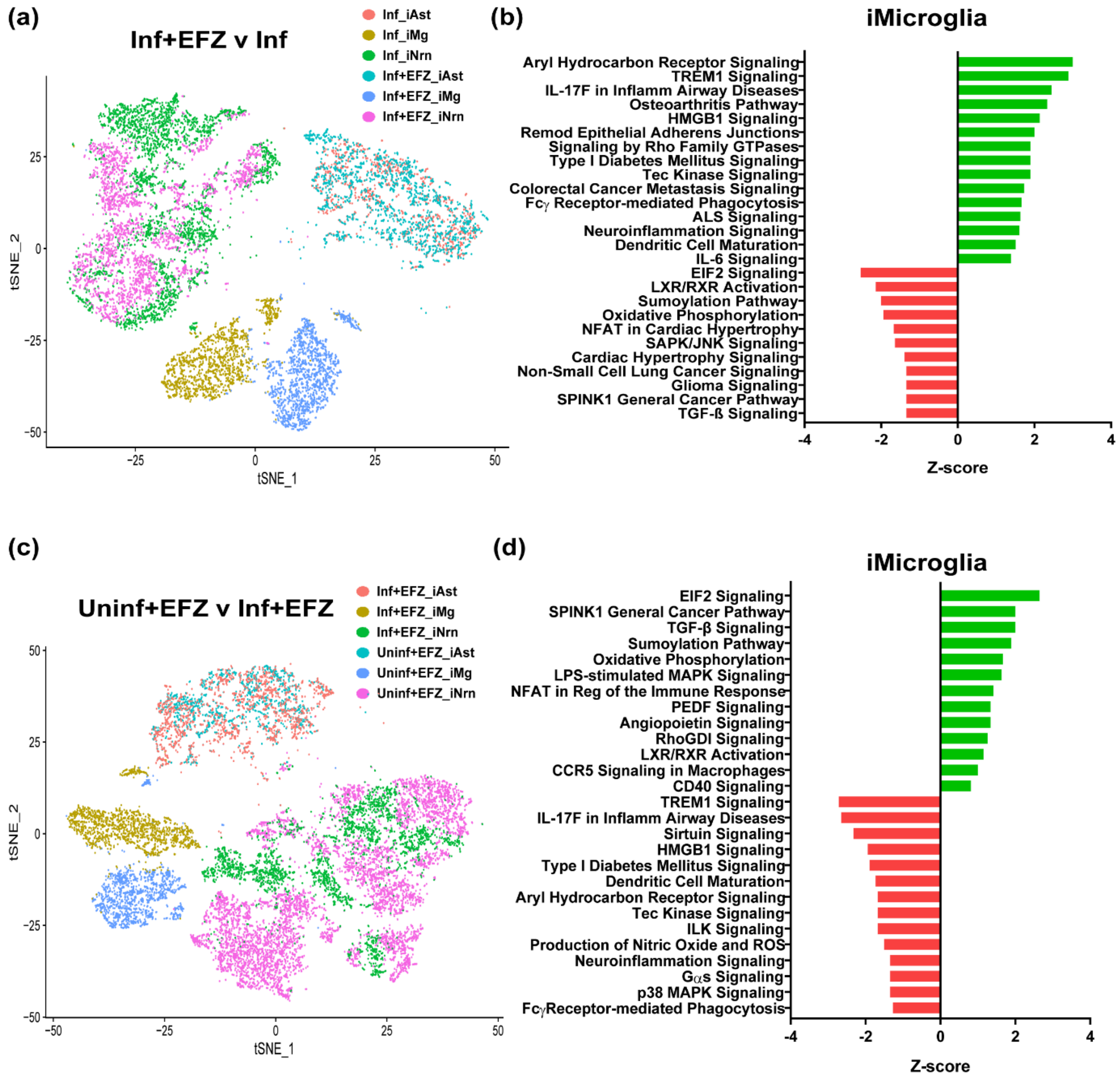


(e)

| Pathway | Genes Dysregulated |
|---|---|
| RhoGDI Signaling | *SRC*, *RHOQ*, *ACTR3*, *RND3*, *CD44*, *ARHGDIA*, *ACTG1*, *MYL12A*, *GNB4*, *PPP1R12A*, *ARPC3* |
| CD40 Signaling | *TANK*, *ICAM1*, *NFKBIA*, *TNFAIP3*, *NFKB1*, *FOS* |

**Figure 6-figure supplement 2: Inf+EFZ treatment yields a distinct inflammatory response in iMg compared to Inf and Uninf+EFZ.**

(a) t-SNE plot of Inf+EFZ and infected conditions. (b) Ingenuity pathway analysis of iMg between Inf+EFZ and infected conditions. Inf+EFZ is baseline. Benjamini-Hochberg FDR 0.05, Fisher's exact <0.05, z-score cutoff ± 0.5. (c) t-SNE plot of Uninf+EFZ and Inf+EFZ conditions. (d) Ingenuity pathway analysis of iMg between Uninf+EFZ and Inf+EFZ conditions. Uninf+EFZ is baseline. Benjamini-Hochberg FDR 0.05, Fisher's exact <0.05, z-score cutoff ± 0.5. (e) Specific genes dysregulated that are involved in the RhoGDI and CD40 pathways in Inf+EFZ iMg compared to Uninf+EFZ iMg. Red genes are downregulated. Green genes are upregulated.

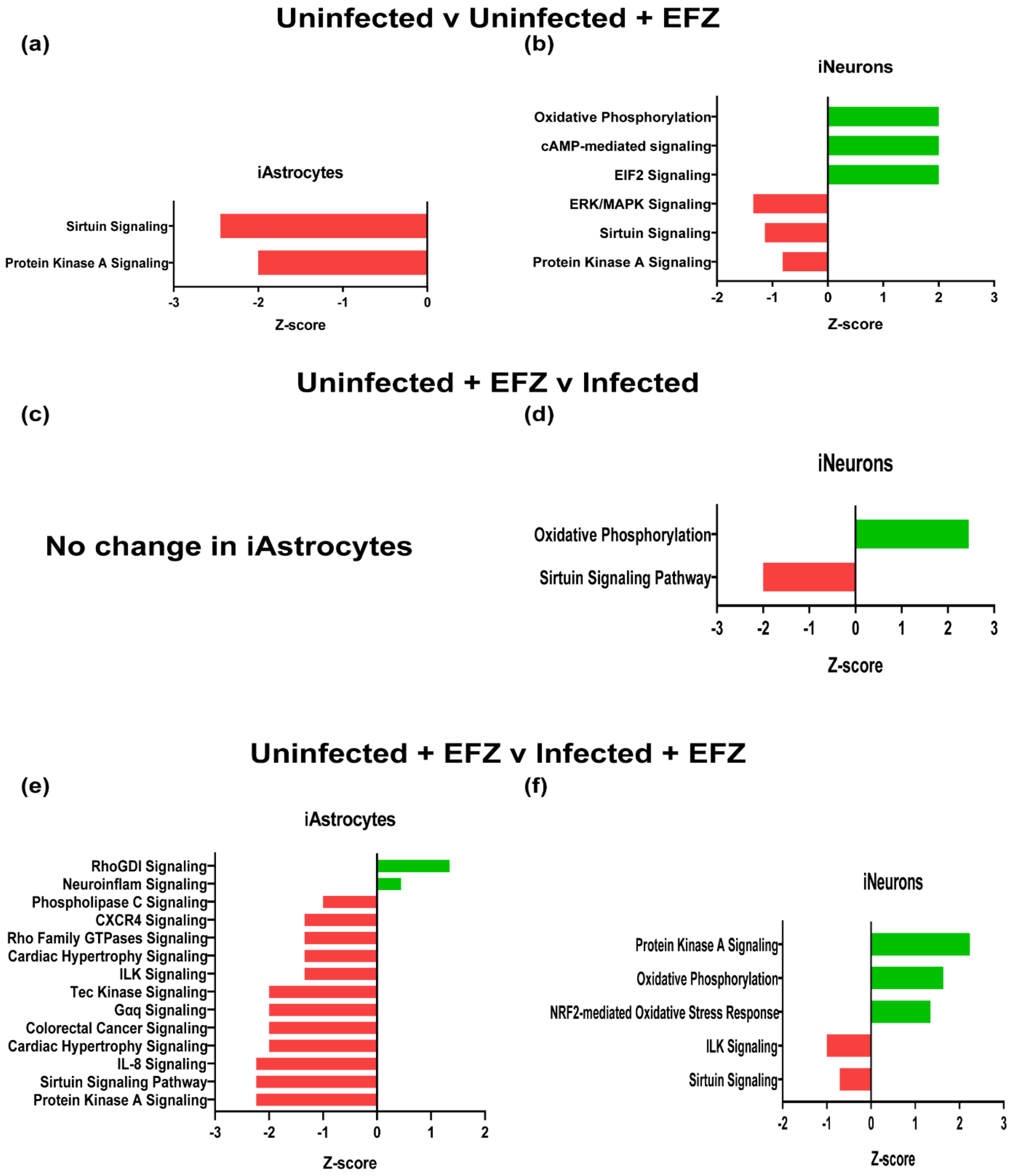


**Figure 7-figure supplement 1: iAst and iNrn top canonical pathways changed in Uninf v Uninf+EFZ, Uninf+EFZ v Inf, and Uninf+EFZ v Inf+EFZ**

(a&b) Top affected pathways from Ingenuity pathway analysis of iAst (a) and iNrn (b) in the uninfected versus uninfected + EFZ conditions. Uninfected is baseline. Benjamini-Hochberg FDR 0.05, Fisher's exact <0.05, z-score cutoff ± 0.5. (c&d) Top affected pathways from Ingenuity pathway analysis of iAst (c) and iNrn (d) in the uninfected + EFZ versus infected conditions. Uninfected + EFZ is baseline. Benjamini-Hochberg FDR 0.05, Fisher's exact <0.05, z-score cutoff ± 0.5. (e&f) Top affected pathways from Ingenuity pathway analysis of iAst (e) and iNrn (f) in the Uninfected + EFZ versus infected + EFZ conditions. Uninfected + EFZ is baseline. Benjamini-Hochberg FDR 0.05, Fisher's exact <0.05, z-score cutoff ± 0.5.

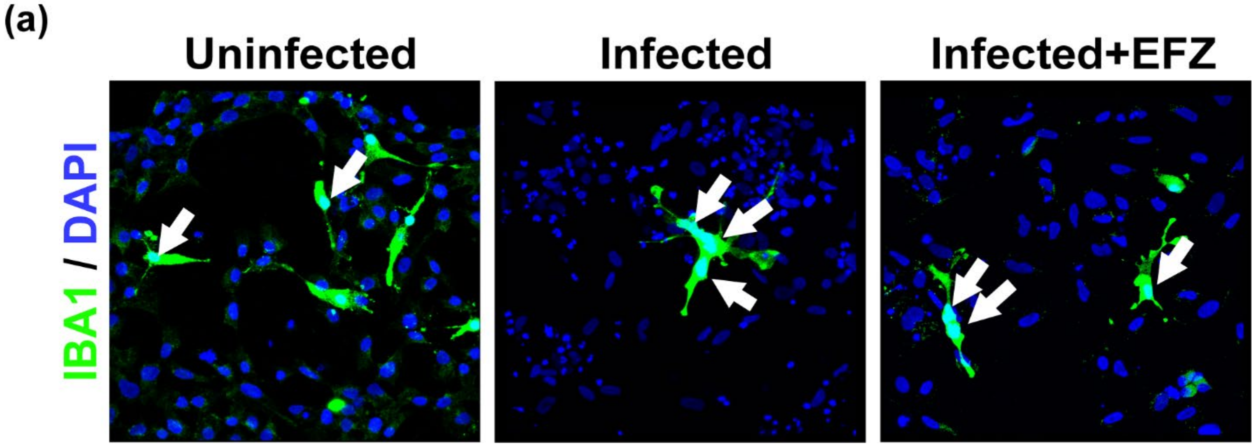


**Figure 8-figure supplement 1: Characterization of tri-culture by scRNA and immunostaining**
(a) Immunostaining of IBA1+ (green) iMg in Uninf, Inf, and Inf+EFZ tri-cultures depicting multi-nucleated iMg in the infected condition and multi- and single-nucleated iMg in the Inf+EFZ condition.

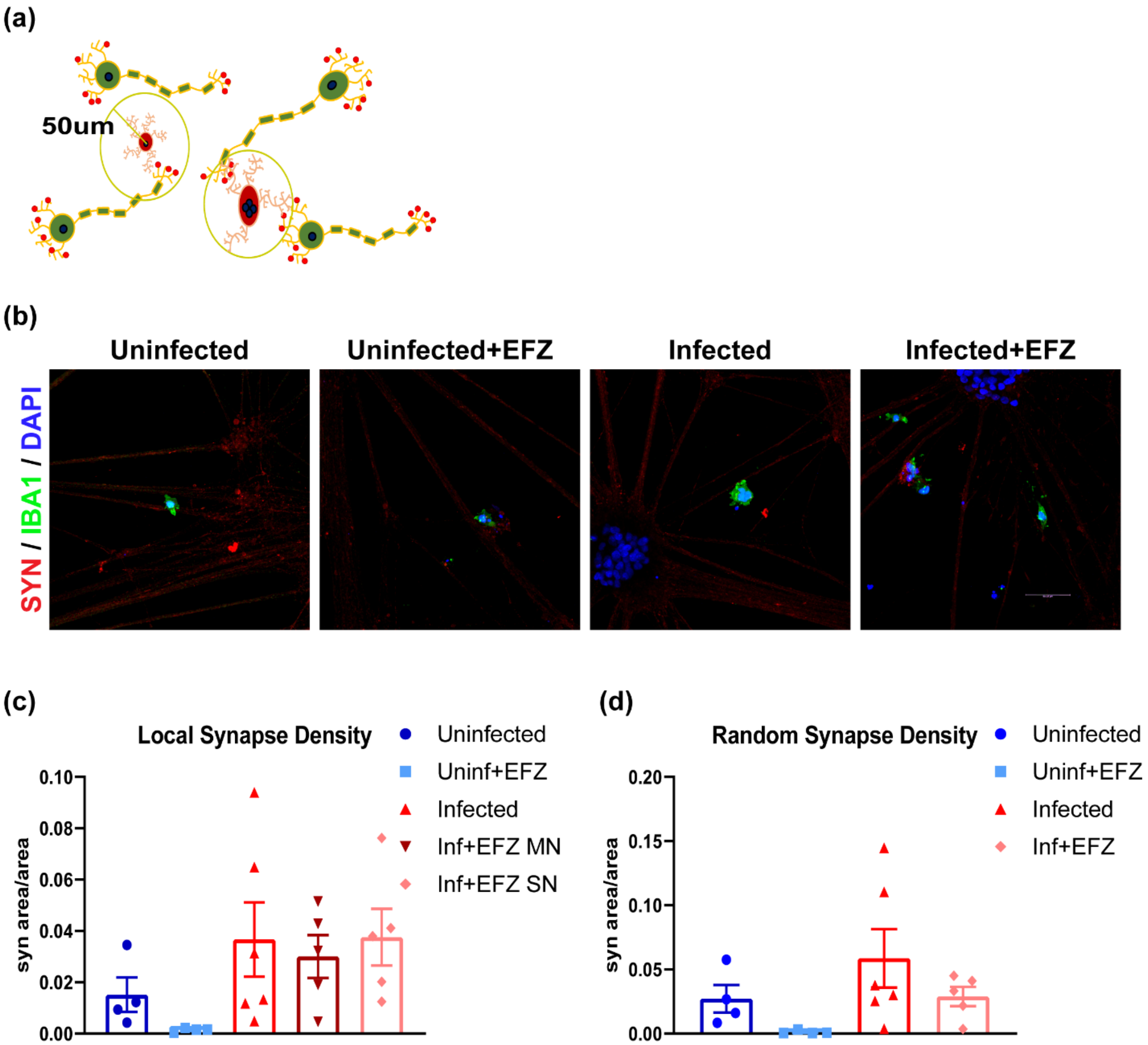


**Figure 8-figure supplement 2: No change in synaptic density local to microglia or throughout the culture**

(a) Cartoon representation and representative images (b) for analysis of local synapse density measured by density of synaptophysin+ (red) staining within a 50um radius of IBA1+ (green) iMg. Scale bar represents 50 µm. (c) No significant difference in local synapse density across conditions compared to control. One-way ANOVA, Dunnett's post hoc analysis, error bars represent SEM. (d) No significant difference in random synapse density across conditions compared to control. One-way ANOVA, Dunnett's post hoc analysis, error bars represent SEM.
